# *In vitro* formation and extended culture of highly metabolically active and contractile tissues

**DOI:** 10.1101/2023.07.07.548141

**Authors:** Isabella A. Bagdasarian, Thamidul Islam Tonmoy, B. Hyle Park, Joshua T. Morgan

**Author notes:** Address correspondence to: Joshua Morgan 900 University Ave Riverside, CA 92521. **Abbreviations:** PDMS; TGF-β2; S-PDMS; P-PDMS; VBAM; BAM; OCT; PS-OCT; PEI/GA – to be filled in if required.

## Abstract

3D cell culture models have gained popularity in recent years as an alternative to animal and 2D cell culture models for pharmaceutical testing and disease modeling. Polydimethylsiloxane (PDMS) is a cost-effective and accessible molding material for 3D cultures; however, routine PDMS molding may not be appropriate for extended culture of contractile and metabolically active tissues. Failures can include loss of culture adhesion to the PDMS mold and limited culture surfaces for nutrient and waste diffusion. In this study, we evaluated PDMS molding materials and surface treatments for highly contractile and metabolically active 3D cell cultures. PDMS functionalized with polydopamine allowed for extended culture duration (14.8 ± 3.97 days) when compared to polyethylamine/glutaraldehyde functionalization (6.94 ± 2.74 days); Additionally, porous PDMS extended culture duration (16.7 ± 3.51 days) compared to smooth PDMS (6.33 ± 2.05 days) after treatment with TGF-β2 to increase culture contraction. Porous PDMS additionally allowed for large (13 mm tall × 8 mm diameter) constructs to be fed by diffusion through the mold, resulting in increased cell density (0.0210 ± 0.0049 mean nuclear fraction) compared to controls (0.0045 ± 0.0016 mean nuclear fraction). As a practical demonstration of the flexibility of porous PDMS, we engineered a vascular bioartificial muscle model (VBAM) and demonstrated extended culture of VBAMs anchored with porous PDMS posts. Using this model, we assessed the effect of feeding frequency on VBAM cellularity. Feeding 3×/week significantly increased nuclear fraction at multiple tissue depths relative to 2×/day. VBAM maturation was similarly improved in 3×/week feeding as measured by nuclear alignment (23.49° ± 3.644) and nuclear aspect ratio (2.274 ± 0.0643) relative to 2x/day (35.93° ± 2.942) and (1.371 ± 0.1127), respectively. The described techniques are designed to be simple and easy to implement with minimal training or expense, improving access to dense and/or metabolically active 3D cell culture models.

## Introduction

Animal models have long been mainstays of biomedical research. While *in vivo* model systems by their nature include important systemic factors, they often fail to recapitulate the physiology of human tissues (Greek and Menache, 2013; Sutherland *et al*., 2013). There is variability in how animals develop pathologies and respond to pharmaceuticals, compared to humans (Lipsky and Sharp, 2001; Dyson and Singer, 2009; Bédard *et al*., 2020). For example, the cholesterol lowering drug cerivastatin was withdrawn from the US market in 2001 after cases of fatal rhabdomyolysis were reported, despite minimal adverse effects being observed in preclinical animal studies (Bischoff and Heller, 1998; Bischoff *et al*., 1998; Keutz and Schlüter, 1998; Thompson *et al*., 2006; Gaukler *et al*., 2016). Two-dimensional (2D) *in vitro* cell culture systems are linked to increased throughput, cost-effectiveness, and experimental simplicity (Breslin and O’Driscoll, 2013; Kapałczyńska *et al*., 2018; Langhans, 2018; Jensen and Teng, 2020). Despite these clear advantages, 2D models are also associated with poor drug sensitivity (Imamura *et al*., 2015; Langhans, 2018; Jensen and Teng, 2020), improper cellular organization and morphology (Soares *et al*., 2012), and altered gene and protein expression (Ravi *et al*., 2015; Costa *et al*., 2016; Jensen and Teng, 2020). In response to these limitations and other needs, significant effort has been placed in the development of 3-dimensional (3D) cell culture models as an additional tool. These systems circumvent the systemic variables, timescale, and ethical challenges associated with animal models, while providing increased physiological relevance compared to conventional *in vitro* methods. (Ravi *et al*., 2015; Jensen and Teng, 2020).

3D tissue culture and organoid formation have become increasingly relevant tools to model complex tissues *in vitro* (Hu *et al*., 2018). Despite progress, several engineering challenges remain. Many tissues have dense cell populations, which are difficult to mimic *in vitro*, due to the high contractile loads produced by cells, leading to collapse of a structured tissue (Kosnik *et al*., 2001). In some cases cell-mediated hydrogel contraction is expected and desired. For example, engineered human skin equivalents often rely on fibroblast induced collagen contraction to render a scaffold with similar mechanical properties to native skin and for use in wound healing studies (Asbill *et al*., 2000; Batheja *et al*., 2009; Garrett *et al*., 2017; Zhang *et al*., 2022). Alternatively, collagen contraction assays can be used to gain insight into the mechanobiology of pathologies such as tissue fibrosis or cancer progression (Provenzano *et al*., 2009; Goetz *et al*., 2011; Le *et al*., 2016; Liu *et al*., 2017; Zhang *et al*., 2022). Despite the evident utility of 3D collagen contraction at times, its uncontrolled occurrence yields highly variable and unreproducible cultures (Machour *et al*., 2022). Specifically, it is well documented that 3D culture collapse alters cell density and hydrogel porosity, due to the reduced volume of the culture (Ahearne, 2014; Machour *et al*., 2022). Ultimately, this compromises the intended geometry and microarchitecture of the culture, and limits overall maturation of the tissue (Gillette *et al*., 2008; Ahearne, 2014; Machour *et al*., 2022). Uncontrolled matrix contraction becomes especially problematic when fabricating naturally contractile cultures, including striated muscle (e.g. skeletal or cardiac muscle) and fibrosis models.

*In vitro* engineered tissues and 3D cell cultures are often formed within a mold, such as polydimethylsiloxane (PDMS). PDMS is a favorable molding material due to its accessibility, biocompatibility, low-cost, optical transparency, and tunable mechanical properties (Brown *et al*., 2005; Chuah *et al*., 2015). Yet, the innate hydrophobicity of PDMS does not promote long term adhesion of hydrophilic extracellular matrix proteins and cells, necessitating the need for separate functionalization steps. Immobilization of 3D cultures to a PDMS mold is frequently accomplished through chemical modification of the surface. One of the most common methods of chemical functionalization is polyethylenemine-glutaraldehyde (PEI/GA) crosslinking of collagen to a PDMS mold, although several other techniques exist for varying biomaterials (Cross *et al*., 2010). Importantly, glutaraldehyde surface treatments are associated with potential adverse health risks (Beauchamp *et al*., 1992; Rideout *et al*., 2005; Zeiger *et al*., 2005) and negative environmental impacts (Leung, 2001; Jolibois *et al*., 2002). Further, PEI/GA treatments are not always successful in immobilizing cell-laden collagen gels during extended culture with high cell densities (Morgan *et al*., 2019).

Polydopamine (PDA), a bioinspired coating agent, is a potential alternative to PEI/GA. A wide variety of material surfaces (including PDMS) can be made adhesive to a broad range of biomolecules via simple dip-coating into an aqueous PDA solution (Lee *et al*., 2007). Importantly, this coating technique does not require specialized equipment, making it accessible to a broad group of researchers. PDA surface coatings of PDMS have been demonstrated to improve adhesion of 2D cell cultures and ECM proteins, such as collagen I (Chuah *et al*., 2015, 2015; Fu *et al*., 2016; Jeong *et al*., 2016; Lee *et al*., 2018; Harati *et al*., 2022). However, it has not yet been demonstrated that PDMS coated with PDA improves 3D cell culture matrix adherence.

Surface roughness of PDMS varies with the fabrication and molding methods, but is generally in the range of 1-20 nm (Prasad *et al*., 2010; Juárez-Moreno *et al*., 2015; Hwang *et al*., 2016; Jang *et al*., 2019). Limited PDMS surface roughness contributes to poor collagen adherence, resulting in hydrogel collapse under cell-generated loads. It has recently been demonstrated that increasing the surface roughness of the PDMS bulk improves collagen film adhesion (Juárez-Moreno *et al*., 2015). Porous PDMS (P-PDMS) is known to have increased surface roughness and can be readily fabricated through the incorporation of sacrificial structures, such as water, salt, or sugar. Importantly, P-PDMS scaffolds can improve biomolecule and cell adhesion relative to normally fabricated smooth PDMS (S-PDMS) (Pedraza *et al*., 2013; Li *et al*., 2018; Jang *et al*., 2019; Riesco *et al*., 2019; Varshney *et al*., 2019). Improved anchoring may be especially beneficial for 3D cultures.

Indeed, strong anchoring of hydrogels to matrix attachment points is especially relevant for engineering contractile tissues, such as skeletal muscle. Tissue engineered skeletal muscle constructs, termed bioartificial muscle models (BAMs), are fabricated from undifferentiated muscle cells (myoblasts) suspended in an extracellular matrix and cast around anchor points within a simple cylindrical mold (Vandenburgh *et al*., 1988). These anchors maintain passive tension within the differentiating tissue. In recent years, the matrix anchoring points have been fabricated from a variety of different materials, including: S-PDMS posts (Vandenburgh *et al*., 2008; Bian and Bursac, 2009; Iuliano *et al*., 2020), 3D printed plastics (Capel *et al*., 2019) and hydrogels (Christensen *et al*., 2020), mesh (Mudera *et al*., 2010; Wragg *et al*., 2019), velcro (Gefen *et al*., 2008; Hinds *et al*., 2011; van der Schaft *et al*., 2013), and silk sutures (Dennis and Kosnik, 2000; Huang *et al*., 2005). Although many of these systems are low-cost and simple to fabricate, there have been reports of the engineered muscle rupturing off the anchors, especially when seeded at higher densities (Smith *et al*., 2012; Wragg *et al*., 2019; Alave Reyes-Furrer *et al*., 2021). P-PDMS may mitigate incidences of tissue rupture due to its increased surface area, allowing for extended culture and maturation of the constructs.

In addition to collagen detachment, 3D cultures also exhibit increased metabolic demands. A well known limitation of 3D cultures and tissue is the diffusion limit; that is, cells do not receive adequate nutrient delivery or waste clearance beyond a few hundred microns of thickness in static culture conditions (Karande *et al*., 2004; Griffith and Swartz, 2006; Buchwald, 2009; Zohar *et al*., 2018; Enrico *et al*., 2022). These challenges hinder the scalability and duration these tissues can be cultured *in vitro*. Efforts to support the metabolic activity of *in vitro* tissues often focus on the development of optimized culture media blends (van der Valk *et al*., 2010; Quiroga-Campano *et al*., 2018); media supplements such as insulin, transferrin, selenium, glucocorticoids, vitamins, and pH buffers are known to improve cell growth and metabolism (van der Valk *et al*., 2010). Further, optimization of feeding may improve cell and tissue health in 3D culture. Often, this is accomplished through the addition of direct tissue perfusion systems designed to increase mass transport within the culture. Perfusion systems have been incorporated in engineered bone (Bancroft *et al*., 2002; Sikavitsas *et al*., 2005; Grayson *et al*., 2008), cartilage (Pazzano *et al*., 2000; Davisson *et al*., 2002), cardiac (Vollert *et al*., 2013), and skeletal muscle (Kim *et al*., 2022) and have been thoroughly reviewed (Martin *et al*., 2004; Li and Cui, 2014; Gelinsky *et al*., 2015). Despite progress, perfusion systems are often optimized for specific research needs and are difficult for non-specialist research groups to utilize. P-PDMS can be fabricated to have interconnected pores (Thurgood *et al*., 2017; Yu *et al*., 2017; Zhang *et al*., 2019). With interconnected pores, there is the potential for P-PDMS to allow for media diffusion through the adhesion surface of the mold.

In this study, we validate the use of P-PDMS as a molding material to support contractile and metabolically active cultures in 3D collagen. Specifically, we compare PEI/GA and PDA chemistries for their ability to stably support contractile cells at high density. Further, we demonstrate that culture media can diffuse through the P-PDMS mold materials, improving health of metabolically active 3D cultures. As a further test case, we demonstrate P-PDMS as suitable anchor points for long term culture of vascularized BAMs (VBAMs); and demonstrate the effect of feeding frequency on skeletal muscle maturation over 5 weeks. Overall, we demonstrate PDA treated P-PDMS molds as a simple and adaptable strategy in 3D cultures, especially where PEI/GA treated S-PDMS molding is unsuitable due to contractile or metabolic concerns.

## Materials and Methods

### Collagen isolation

Collagen Type I was isolated from rat tail tendons as previously described (Rajan *et al*., 2006; Cross *et al*., 2010; García-Gareta, 2014). Briefly, collagen fibers were retrieved from the fibers of rat tail (Pel-Freez Biologicals, Rogers, AR) tendons and soaked in 1x PBS. Afterwards, fibers were incubated in acetone and 70% isopropanol for 5 min each. Fibers were split evenly among conical tubes and swelled in 0.1% glacial acetic acid for 7 d on a rocker at 4°C. Dissolved collagen was centrifuged at ∼20,000 x g for 1 h at 4°C to remove impurities. The collagen-containing supernatant was frozen at -80°C overnight and lyophilized to generate a collagen sponge. Prior to use, collagen sponges were dissolved in 0.1% glacial acetic acid to 8 mg/mL and stored at 4°C.

### Cell culture

All cells were routinely cultured at 37°C and 5% CO2. IMR90s (human lung myofibroblast; passage 17-19; ATCC, VA, USA), MDCKs (canine kidney epithelial; ATCC), HMEC1s (human microvascular endothelial; passage 8-10; ATCC), C2C12s (mouse myoblasts; passage 19; ATCC), and ASC52telos (human adipose derived stem cells; passage 7, ATCC), were maintained in their respective media blends (**Table 1**) prior to 3D culture.

Fluorescent expressing C2C12s and HMEC1 cells were generated using lentiviral transduction. Transgenes were transduced into cells through 2^nd^ generation lentiviral system. Briefly, 70% confluent HEK293TN (System Biosciences, Palo Alto, CA) cells were triple transfected (TransIT-Lenti, Mirus Bio, Madison, WI) with plasmids for viral packaging (psPAX2 was a gift from Didier Trono; Addgene plasmid #12260; http://n2t.net/addgene:12260; RRID:Addgene_12260), viral envelope (pMD2.G was a gift from Didier Trono; Addgene plasmid #12259; http://n2t.net/addgene:12259; RRID:Addgene_12259). At 48 and 72 h post transfection, viral supernatant was collected, centrifuged to remove cellular debris, and filtered through a 0.45 μm cellulose acetate filter (Corning). Viral particles were stored at 4°C and used within 48 h by addition to culture media at a 1:2 ratio. Fluorescent expressing C2C12s were created by introducing copGFP (pCDH-EF1-copGFP-T2A-Puro was a gift from Kazuhiro Oka (Addgene plasmid # 72263; http://n2t.net/addgene:72263; RRID:Addgene_72263) and fluorescent HMEC1s were created by introducing mCherry (pCDH-CMV-mCherry-T2A-Puro was a gift from Kazuhiro Oka (Addgene plasmid # 72264; http://n2t.net/addgene:72264; RRID:Addgene_72264) plasmids. Positively expressing cells were selected using Puromycin at 1 µg/mL (C2C12 cells) or 0.1 µg/mL (HMEC1 cells) for 2 weeks.

### S-PDMS mold fabrication & surface treatments

PDMS (Sylgard 184; Dow Corning, Midland MI) prepolymer was molded around a 3D printed mold (acrylonitrile butadiene styrene) creating wells 7 mm in diameter and 1.5 mm deep (**Figure 1B**). S-PDMS was cured in the oven for 48 h at 55 °C and autoclaved prior to functionalization with (poly)ethylenimine/glutaraldehyde (PEI/GA) or polydopamine (PDA). Briefly, naïve S-PDMS molds were incubated in 2% PEI for 30 min, rinsed 3 times with autoclaved deionized water, dried, and immersed in 0.2% GA for 1 h, rinsed, and dried again (Cross *et al*., 2010; Morgan *et al*., 2019). Alternatively, naïve S-PDMS molds were incubated overnight in 2 mg/mL PDA solution made from dopamine hydrochloride (Sigma-Aldrich, St. Louis, MO) in 10 mM Tris Buffer (pH ∼8.5; Apex Bioresearch Products, Boston, MA) as previously described (Chuah *et al*., 2015). S-PDMS molds were rinsed in autoclaved deionized water and dried prior to cell culture.

**Figure 1:**
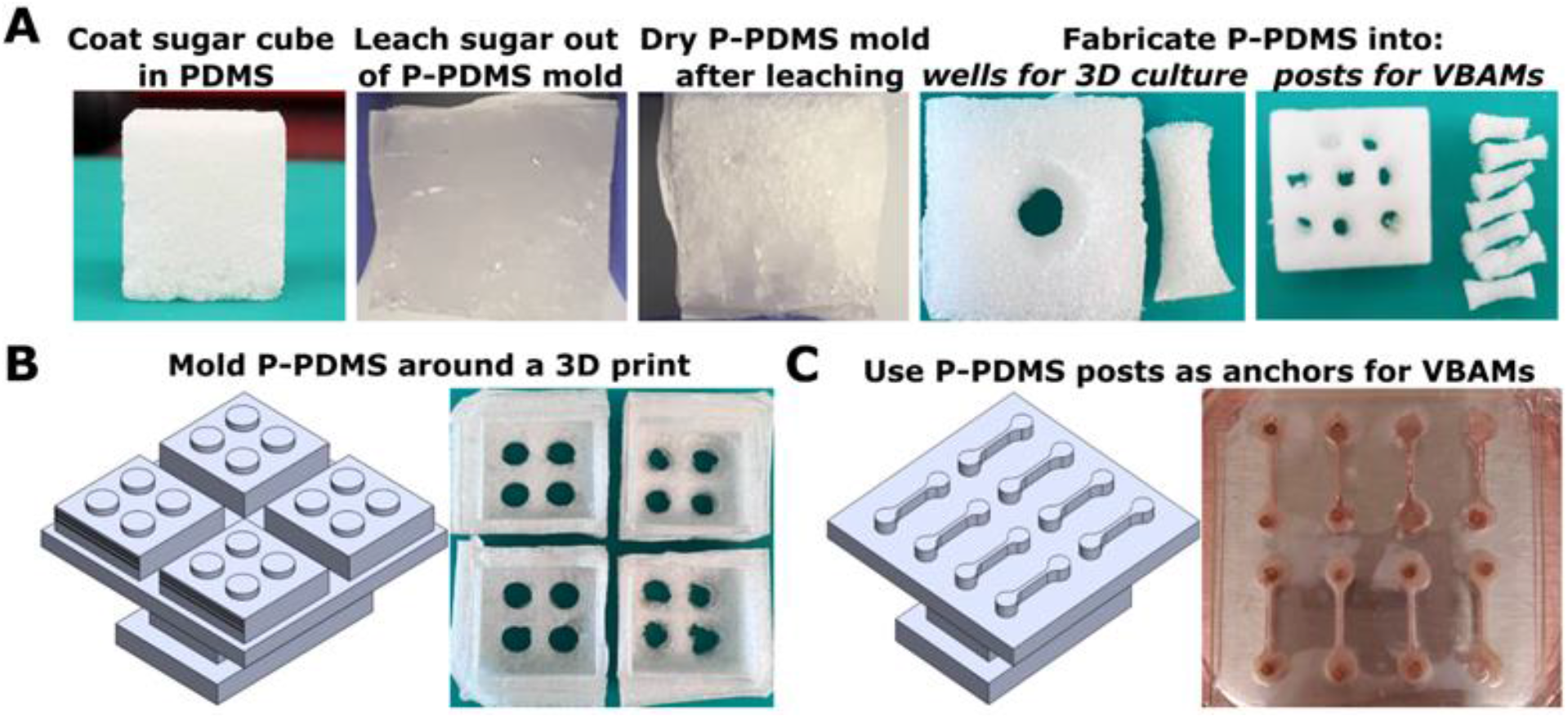
Fabrication of S-PDMS and P-PDMS molds. (A) PDMS pre-polymer is cast around a sugar cube template and thermally cured. Sugar granules are leached out in water and the molds are dried, revealing P-PDMS. (B) P-PDMS is molded around a 3D print to form wells 7 mm in diameter and 1.5 mm tall. (C) S-PDMS is molded around a 3D print to form VBAM outer chambers (20 mm x 1 mm x 1.5 mm). P-PDMS posts are punched to be 1/8" in diameter and adhered to glass inside the S-PDMS outer chambers.

### P-PDMS mold fabrication

Two methods were used to form P-PDMS molds for 3D cell culture using sugar as a porogen. The first method, used in the collagen contractility study, consisted of packing 20 g granulated sugar into a 90 mm petri dish around the 3D printed mold. 15 g of PDMS prepolymer was poured onto the sugar and around the mold and cured for 48 h at 55°C. The P-PDMS was demolded from the 3D print and leached in water for 7 d at 40° C (**Figure 1A**). To verify sugar had been fully removed and to dry the P-PDMS, molds were placed in the oven at 120°C for 2 h. If browning was observed, this was taken as an indication of sugar caramelization indicating incomplete porogen removal; these molds were discarded. P-PDMS molds were adhered to 22 mm x 40 mm glass coverslips with a thin layer of PDMS prepolymer, cured in the oven, and autoclaved prior to functionalization as described above. For 3D culture metabolic studies, P-PDMS was fabricated using sugar cubes as a sacrificial template to ensure pores were interconnected (Ren *et al*., 2017; González-Rivera *et al*., 2018; Li *et al*., 2018). Sugar cubes were coated in PDMS prepolymer under vacuum, and cured at 55°C as described, followed by sugar dissolution in the water bath for 7 days. 3D culture molds were formed using a 1/4” punch to create a well for collagen cultures, cut to be ½” tall, then adhered to glass coverslips (22 mm x 40 mm) with PDMS prepolymer (**Figure 1A**). Prior to PDA functionalization, P-PDMS was rewet in an ethanol gradient. Molds were submerged in 100% EtOH for 2 hrs, followed by overnight incubation in 50% EtOH:H_2_O, 1 hr in 25% EtOH:H_2_O, and 1 hr in 100% H_2_O the following morning. Infiltration of water into the pores was readily visible when compared to dried molds (**Figure 1A**). Porosity was further confirmed by loading the center well and observing leakage to the outside. VBAM mold posts were also fabricated from P-PDMS sugar cubes. Posts for matrix attachment were made using a 1/8” diameter punch and cut to be ∼3 mm tall. Posts were functionalized in PDA, dried, and were adhered to glass as described above. S-PDMS outer chambers were fabricated from PDMS prepolymer molded around a 3D print design (acrylonitrile butadiene styrene). Each outer chamber has dimensions of 20 mm x 1 mm x 2.5 mm (LxWxH) (**Figure 1C**). After curing, S-PDMS was demolded and individual chambers were trimmed, placed around the posts, and autoclaved.

### Assessment of PDMS surface treatments

S-PDMS molds were prepared and treated with either PEI/GA, PDA, or left naïve (control) as described. IMR90s were suspended in 3 mg/mL rat tail collagen type I at 2 x 10^6^ cells/mL and gelled at 37°C for 30 min and then submerged in IMR90 medium (**Table 1**). Cultures were monitored daily for loss of adhesion from the S-PDMS mold. Cultures were considered de-adhered when more than 270° of the perimeter had detached from the S-PDMS wall. Survival past this point was quantified relative to the number of days it took the naïve cultures to detach (**Figure 2**).

**Figure 2:**
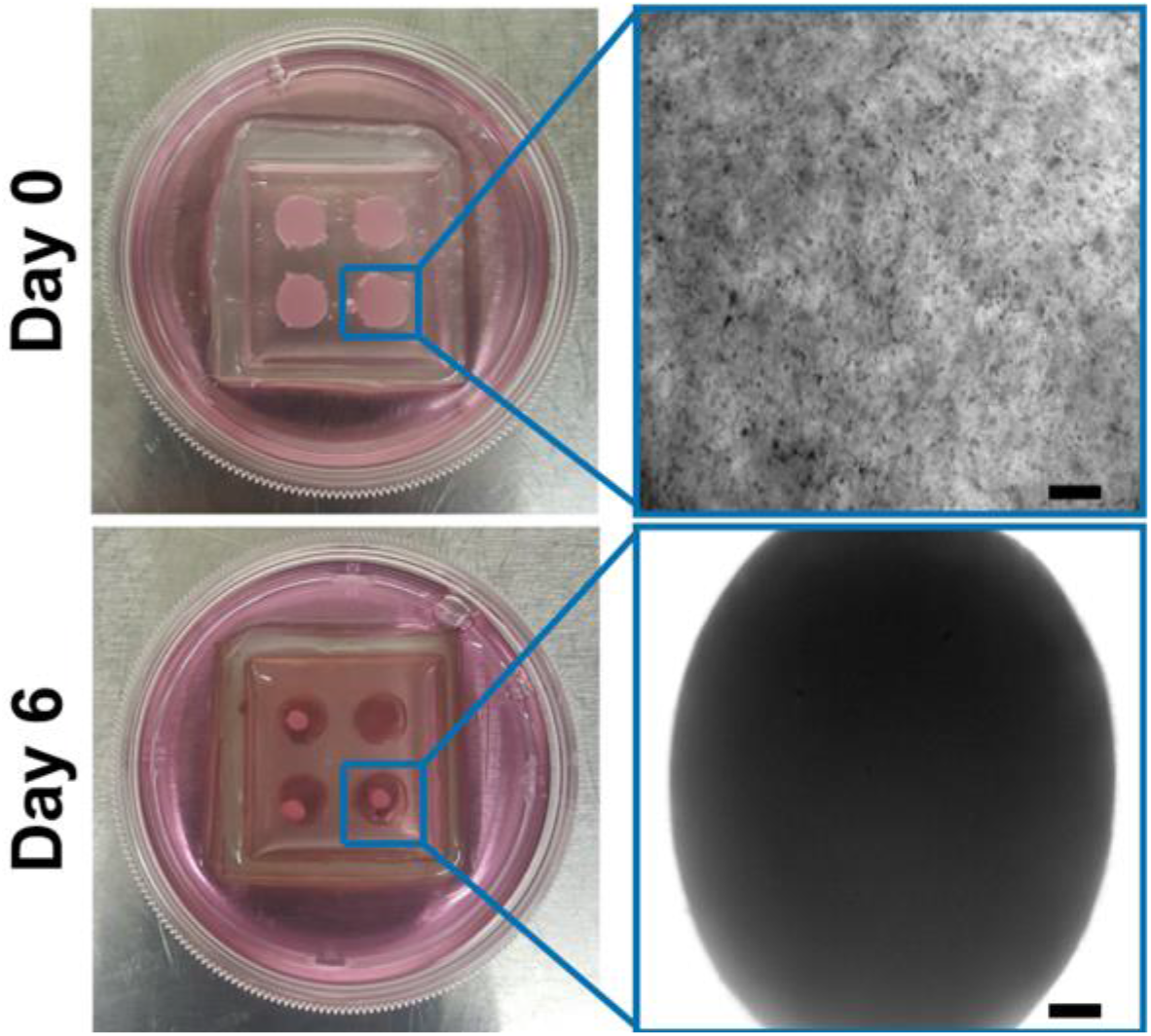
Loss of S-PDMS/collagen adhesion. (Top) IMR90s are seeded in a collagen matrix in a naïve S-PDMS mold. (Bottom) By day 6 of culture the collagen has completely detached from S-PDMS wall.

### Assessment of P-PDMS/collagen adhesion

S-PDMS and P-PDMS molds were prepared, treated with PDA, and seeded with IMR90s at 2 x 10^6^ cells/mL cells/mL as described above. After gelation samples were submerged in IMR90 medium with/without 2 ng/mL TGF-β2 (100-35B; PeproTech, Cranbury, NJ). Cultures were monitored for loss of collagen adhesion and contraction of the gel for 21 d and survival was quantified as described above.

### Assessment of P-PDMS nutrient diffusion

For metabolic studies, 13 mm tall × 8 mm diameter S-PDMS and P-PDMS molds were fabricated and seeded with MDCK cells at 2 x 10^6^ cells/mL. In the case of the P-PDMS molds, collagen that leaked through the porous mold was cut away after gelation. The molds and collagen were placed in 12 well plates and media was added to be level with the mold surface, but not covering the culture (the top of the culture was at the air-liquid interface). Samples were fed at days 1,2, and 4 of culture. Media was isolated for downstream glucose assays at days 1 and 5. After 5 days of culture, samples were fixed for immunofluorescence.

### VBAM model fabrication

To facilitate live imaging throughout culture, VBAMs were fabricated from C2C12s expressing copGFP and HMEC1 cells expressing mCherry. C2C12s, HMEC1s, and ASC52telos were trypsinized and suspended in a 2 mg/mL type I rat tail collagen matrix (3440-100-01; R&D Biosystems, Minneapolis, MN) at 10 x 10^6^ cells/mL, 2 x 10^6^ cells/mL, and 0.5 x 10^6^ cells/mL, respectively. Cell laden collagen was seeded in S-PDMS chambers 20 mm x 1 mm x 2.5 mm around P-PDMS posts 1/8” in diameter (**Figure 1C**). After a 60 minute gelation, constructs were submerged in VBAM growth media (**Table 1**) supplemented with 1 ng/mL VEGF. VBAM growth media was exchanged every other day until day 5 of culture to allow for myoblast proliferation. At this time media was replaced with VBAM differentiation media (**Table 1**). Microvessel self-assembly was promoted through VEGF supplementation for another 2 weeks before being replace with 0.1 ng/mL PDGF-BB for the remainder of the culture period (Chen *et al*., 2007; Han *et al*., 2013). These conditions were shown to result in robust microvessel networks in prior experiments without muscle cells (**Figure S1**). After 3-4 days in differentiation media, S-PDMS chambers were removed to expose a larger surface area of the tissue to culture media. VBAMs were differentiated for 5 weeks prior to tissue fixation.

To test the effects of sample feeding frequency during differentiation we implemented the following media exchange regimes: 3x a week on Monday, Wednesday, and Friday (MWF), daily, or every 12 h (B.I.D). Feedings performed every 12 hrs were controlled via an automated syringe pump system. Briefly, differentiation media was prepared in an autoclaved beaker in parallel with the MWF differentiation media. Prior to feeding, 2 pieces of tubing (0.07” outer diameter, 18” long, Cole-Parmer, Vernon Hills, IL) and 2 18 ga. steel blunt tip dispensing cannulas were sterilized in autoclave pouches. 90 mm tissue culture dish lids were drilled with a 1/16” bit to create 2 holes on opposite ends for tubing insertion. Lids were sterilized in 70% ethanol prior to use. For media infusion a sterile 60 mL plastic Luer slip syringe was filled with 60 mL VBAM differentiation media and connected to an 18 ga. steel dispensing tip and one of the tubing sections. Tubing was primed with media and insert into the tissue culture dish lid and stabilized with tape. Similarly, a media withdrawal syringe was connected to the remaining dispensing tip and tubing section and insert into the dish lid, ensuring that the tubing end was flush with the bottom of the plate. The plate was primed with 13 mL of media. The media exchange system and VBAM samples were then transferred to the cell culture incubator and tubing was taped in place. Each syringe was connected to a multi-step programmable syringe pump (Chemyx; Stafford, TX). Media withdrawal occurred every 12 h and media infusion occurred every 12 hr with a 15 minute delay to ensure a fresh bolus was delivered.

### 3D culture fixation and staining

3D IMR90 cultures were fixed in 4% paraformaldehyde in PBS with 0.5% Triton X-100 *in situ* for 2 h at room temperature followed by an overnight permeabilization in 0.5% Triton X- 100 in PBS at 4°C. Following fixation, samples were demolded and placed in 1.7 mL tubes and stained against F-actin and nuclei using Phalloidin and DRAQ7 in blocking buffer (**Table 2**) on a rocker for 48 h at 4°C. 3D MDCK cultures were fixed 4% paraformaldehyde in PBS with 0.5% Triton X-100 *in situ* for 2 hr at room temperature followed by an overnight fixation and permeabilization in fresh 4% paraformaldehyde and 0.5% Triton X-100 in PBS at 4°C. After fixation, samples were demolded and transferred to a 48 well plate to maintain spatial orientation during staining. Nuclei were labeled with DRAQ7 as described above.

To ensure sufficient penetration of antibodies into VBAM models, we adopted a modified version of Dent’s fixation (Dent *et al*., 1989; Ahnfelt-Rønne *et al*., 2007). Briefly, VBAMs were triple rinsed in PBS containing calcium and magnesium (21-030-CM; Corning). Samples were fixed *in situ* in [4:1] methanol and DMSO (BP231-1; Fisher BioReagents; Waltham, MA) at 4°C overnight. The following day VBAMs were demolded and transferred to a 24 well plate and dehydrated with 3x methanol incubations for 20 min at 4°C. After, samples were incubated in Dent’s bleach solution consisting of [4:1:1] methanol:DMSO:30% hydrogen peroxide for 2 h at room temperature. Samples were rehydrated in a descending methanol gradient in PBS: 100%, 75%, 50%, 25% methanol and 100% 1× PBS for 10 min at room temperature, before incubating in blocking buffer for 2 h. VBAMs were stained against markers of terminal muscle differentiation and for mCherry labeled HMEC1 cells (**Table 2**) for 72 h on a rocker at 4°C followed by secondary antibody staining for 48 h on a rocker at 4°C (**Table 2**).

### Image acquisition, tissue clearing, and volumetric quantification

All samples were imaged on a Leica TCS SPE-II laser scanning confocal microscope (Leica, Buffalo Grove, IL). For 3D culture experiments with IMR90s, MDCKs, and VBAMs, a total volume of 0.5 mm^3^, 3 - 7.5 mm^3^, and 0.25 - 1.20 mm^3^ were acquired, respectively. Samples were imaged in S-PDMS chambers attached to glass with PBS and acquisition settings were held constant within experimental groups. VBAM and MDCK experiments were imaged before and after clearing. Tissue clearing was performed similar to previously published protocols (Sanchez and Morgan, 2021). Briefly, samples were dehydrated in excess methanol on a rocker at 4°C 3× for 20 min. Samples were then transferred to glass petri dishes and cleared in methyl salicylate for 7× 10 min incubations at room temperature. Samples were imaged in methyl salicylate in S-PDMS chambers sealed to coverslip glass with silicone grease. Confocal tilescans were acquired of the culture base and stitched into a single volume for MDCK experiments. VBAMs were imaged end to end via sequential tilescans.

Prior to analysis all tilescan volumes were aligned and stitched together using a custom MATLAB implementation of the Phase Correlation Method previously described (Preibisch *et al*., 2009). Nuclear volume fraction was quantified using custom algorithms. Briefly, noise was removed from the nuclei channel using a median filter with a 1.07 µm x 1.07 µm x 7.23 µm kernel and background illumination was smoothed via tophat filtering with a sphere structuring element (2.1 µm). Intensities were scaled volumetrically using a linear image adjustment, and additional noise was removed with median filtering using a 1.79 µm x 1.79 µm x 12.0 µm kernel. Nuclei were segmented using hysteresis thresholding with empirically determined limits of 75 and 85. Finally, small artifacts were removed from the binary volume via area opening. Nuclear volume fraction was defined as the sum of nuclei positive voxels in the binary volume and divided by the total voxel number.

To quantify annular nuclear volume fraction in VBAMs, we first stitched and filtered tilescans as described above. Muscle bulk was segmented using the myosin or titin channel via hysteresis thresholding. Debris and artifacts were removed from the binary muscle bulk volume with morphological opening and closing using a disk structuring element of radius 1.79 µm and 5.38 µm, respectively. Similarly, the nuclei channel was segmented via hysteresis thresholding and artifacts were removed. A Euclidean distance transform was performed on the muscle bulk starting at the outer edge of the nuclear layer and used to determine voxel location from the tissue surface. Annular rings moving from the tissue surface inwards were spaced at 14.3 µm. Nuclear volume fraction was defined within each annulus. Annular nuclear fraction was quantified for 5 annular rings, with a max depth of 75 µm (**Figure 3**).

**Figure 3:**
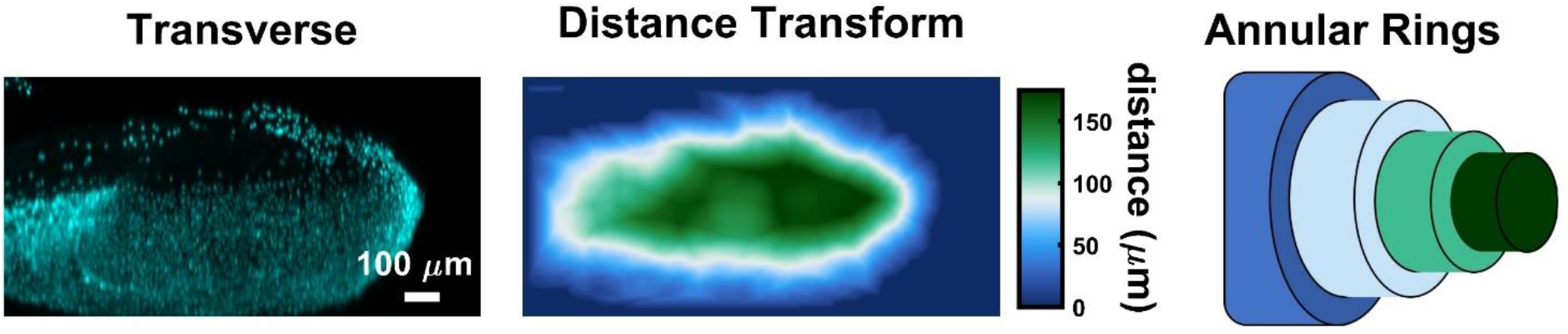
Annular nuclear volume fraction demonstration. VBAM nuclei (transverse) and muscle volumes are segmented. A distance transform is performed on the segmented muscle volume to define how far each pixel in the tissue is from the tissue surface. Annular rings are defined using the distance transform volume and integrated nuclear intensity is quantified within them.

To quantify nuclear angle and aspect ratio we stitched and filtered the VBAM tilescans and segmented the muscle bulk as described above, with an additional watershedding step to separate individual nuclei. Nuclear eigenvectors and principal axis length were output. To account for tilt in the acquired tilescans we solved for local muscle eigenvectors along the skeletonized muscle volume. To accomplish this, we defined seed points at opposite edges of the muscle volume and used an adaption of fast marching skeletonization previously described (Van Uitert and Bitter, 2007; Morgan *et al*., 2019; Shirazi *et al*., 2019). Local 3D orientation along the skeletonization and nuclear angle was quantified as the orientation of the major axis of a best fit ellipsoid. Nuclear aspect ratio was quantified by solving for the ratio of nuclei major principal axis length to minor principal axis length, where a perfectly spherical nuclei would have an aspect ratio equal to 1. Example code used to quantify VBAM annular nuclear volume fraction, nuclear angle, and aspect ratio is provided (**Supplementary Materials**).

### Polarization-sensitive optical coherence tomography

Four fixed and uncleared VBAM samples (two MWF and two B.I.D) were scanned with a custom-built spectral domain PS-OCT system, a detailed description of which is described in a previous report (Wang *et al*., 2012). The system has a center wavelength of 1310 nm with axial and lateral resolution of 11 µm and 37 µm, respectively. The imaging depth is 2 mm in the air with a sensitivity roll-off of 13 dB over that range. The VBAM samples were submerged in 1× PBS during image acquisition to reduce specular reflection from the surface and were pinned at the ends. The sample region between the two pinned ends were scanned volumetrically in overlapping volumes with 1-2 mm overlap between the volumes. The volumes were manually aligned during post-processing. Each volume is comprised of 200 cross-sectional images which encompasses 4.5 mm length of the sample. The cross-sectional images have 1024 A-lines encompassing 1.125 mm width.

The post-processing of the volumetric data was performed with MATLAB. The structural intensity images were generated with standard Fourier domain processing method (Mitsui, 1999; Wojtkowski *et al*., 2002; Choma *et al*., 2003; De Boer *et al*., 2003; Nassif *et al*., 2004) which shows the intensity of light backscattered from the samples (**Figure 9A & 9E**). Muscle tissue also exhibits form birefringence (Haskell *et al*., 1989), an optical property arising from the structural anisotropy of long, parallel fibrils embedded in a medium of different refractive index. When polarized light passes through a birefringent medium, the two orthogonal polarization components of light travel at different speeds due to the difference in their refractive indices introducing a phase retardation between the two components. PS-OCT can measure this phase retardation and optic axis (Hee *et al*., 1992; De Boer *et al*., 1997; Park *et al*., 2001, 2005) which are measures of the degree of organization and orientation of the fibrous structure, respectively. We used the spectral binning algorithm (Villiger *et al*., 2013) to measure the local phase retardation 𝜌 (**Figure 9B & 9F**) and optic axis vector (𝑞, 𝑢, 𝑣). Ideally, the measured optic axis should lie on the QU-plane on a Poincaré sphere representation. However, because of the birefringence of the fiber-based imaging system, the measured optic axis vector is rotated off the QU-plane (Park *et al*., 2005). From the rotated optic axis, we obtained the relative orientation of the muscle fibers by measuring the orientation angle 𝜃 with respect to a manually defined reference angle (**Figure 9C & 9G**).

Nam *et* al. (2018) recently proposed vectorial birefringence, defined as 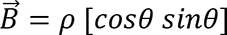, which adds directional contrast to the scalar birefringence used in previous studies (Baumann, 2017; De Boer *et al*., 2017; Nam *et al*., 2018). The vectorial birefringence is presented in HSV colormap where the H channel sets the color and the V channel sets the brightness of the color. In the vectorial birefringence images (**Figure 9D & 9H**), the color encodes the muscle fiber orientation 𝜃 and the brightness of the color scales with the phase retardation 𝜌. The brightness was multiplied by the intensity to exclude noise dominant pixels. The cross-sectional vectorial birefringence images were then compiled in Amira for generating 3D images of the samples (**Figure 9I-J**).

For quantitative comparison, we used the phase retardation of the samples. In each cross-section, the tissue region was first identified using an intensity-based threshold of the background (60 dB). The average phase retardation inside these tissue regions were measured and plotted as a function of length of the samples in **Figure S2A**. The mean phase retardation over the length of the samples were then used for quantitatively comparing the two experimental groups using two sample t-test (**Figure 9K**).

### Glucose assay

A colorimetric glucose assay (10009582; Cayman Chemical, Ann Arbor, MI) was performed according to manufacturer’s directions. Briefly, isolated media samples and control MDCK media were diluted [1:25] in assay buffer. Glucose standards and diluted samples were added to a 96 well plate in duplicate and combined with provided colorimetric enzyme mixture. The plate was incubated for 10 min at 37°C before reading on a plate reader (SpectraMax M2, Molecular Devices, San Jose, CA) at 513 nm. Output absorbances were corrected by taking the mean of each duplicate and subtracting the mean absorbance of the 0 mg/dL glucose standard. Corrected absorbances were normalized to the corrected absorbance of control MDCK media.

### Data Analysis & statistics

For 3D IMR90 experiments, data was reported as mean days without collagen detachment ± SEM. PEI/GA and PDA surface treatments (n = 4) were tested for statistically significant differences using a paired sample t-test. Comparisons of S-PDMS and P-PDMS cultures (n = 3) were tested for statistically significant differences using a 2-way analysis of variance (ANOVA) followed by Tukey’s honestly significant difference (HSD) *post hoc* test for effects of +/- TGF-β2 and S-PDMS vs P-PDMS cultures. Similarly, glucose assay (n = 3) results are reported as mean normalized absorbances and were tested using ANOVA followed by Tukey’s HSD post hoc test. Nuclear fraction quantification is reported as mean ± SEM and assessed using a two-sample t-test For VBAMs each sample fabricated was treated as a biological replicate (n = 6). Phase retardation is reported as mean ± SEM (n = 2) and assessed using a two-sample t-test. Annular nuclear fraction, nuclear angle, and nuclear aspect ratio are reported as mean ± SEM and data was compared with a paired t-test (n = 6).

## Results

### Polydopamine surface treatment extends S-PDMS/collagen adhesion

S-PDMS surfaces were functionalized with PEI/GA or PDA and seeded with IMR90s suspended in a collagen gel, as described above. All samples were fed regularly and were monitored until loss of collagen adhesion. PEI/GA and PDA culture detachment was quantified relative to the detachment of naïve controls (e.g. days past naïve detachment). PEI/GA samples survived on 6.94 ± 2.74 days past naïve, while PDA cultures survived 14.8 ± 3.97 days (**Figure 4A**). To visualize the cell population in PDA samples, we fixed at day 4 and day 12 and stained for f-actin and nuclei. Confocal imaging of these samples revealed high cellularity with spread morphology and stress fibers, consistent with a dense, highly contractile culture (**Figure 4B**).

**Figure 4:**
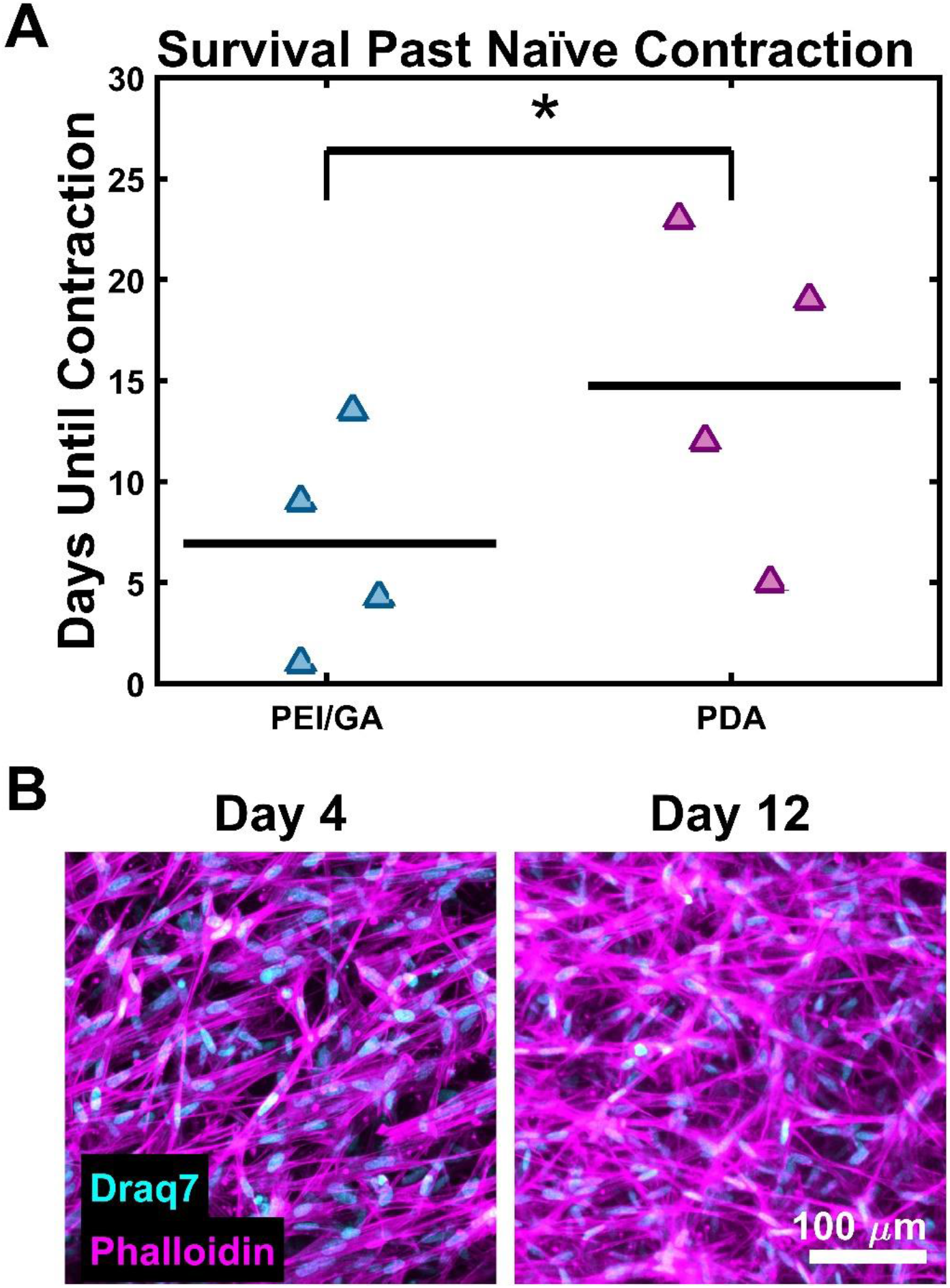
Effect of S-PDMS surface treatment on collagen adhesion. (A) Quantification of S-PDMS/collagen adhesion relative to the naïve control for each replicate reveals that samples in PDA molds maintained adherence longer than PEI-GA molds (n = 4, p < 0.05). Mean is indicated by black bars and triangle markers are data points. (B) PDA cultures were fixed at day 4 and day 12 and imaged for f-actin and nuclei. Representative intensity-based maximum projections qualitatively confirm cultures had high cell density at both time points.

### Porous PDMS extends PDMS/collagen adhesion during contractile culture

S-PDMS and P-PDMS molds were fabricated and treated with PDA prior to seeding with IMR90s in collagen. To increase culture contraction, samples were cultured without/with ((-)/(+)) 2 ng/mL TGF-β2 after day 1 of culture. Samples were fixed and stained against f-actin and nuclei at day 11 of culture to compare cellularity. Samples cultured with TGF-β2 ((+) TGF-β2) demonstrated a qualitative increase in stress fiber formation in both S-PDMS and P-PDMS molds (**Figure 5A & 5B**). There were no qualitative differences in cell density between S-PDMS and P-PDMS samples (-)/(+) TGF-β2. S-PDMS cultures (-) TGF-β had a mean time to detachment of 10.3 ± 4.22 days, the presence of TGF-β2 reduced that to 6.33 ± 2.05 days. P-PDMS cultures (-) TGF-β2 showed no indications of contraction or loss of adhesion with the mold; at day 21, all remaining cultures were ended. With TGF-β2, P-PDMS samples still demonstrated a significant increase in time before detachment when compared to S-PDMS (+) TGF-β2, 16.7 ± 3.51 days.

**Figure 5:**
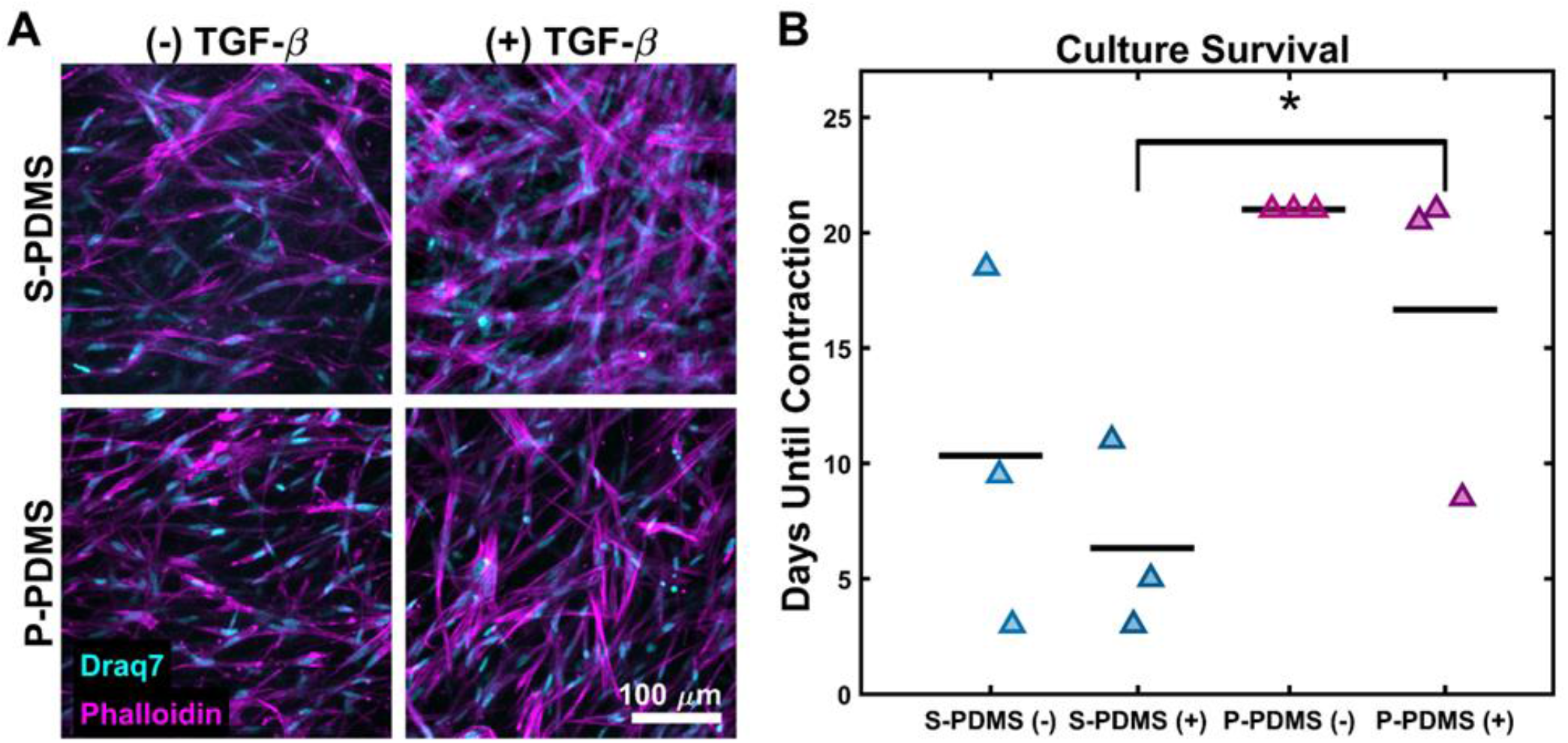
Effects of PDMS surface area on collagen adhesion. (A) IMR90s suspended in collagen gels molded by PDA treated S-PDMS or P-PDMS were fixed and stained for f-actin and nuclei at day 11. Representative intensity-based maximum projections demonstrate consistent cellularity regardless of molding method or TGF-β2 supplementation. (B) Culture survival was defined as the number of days samples lasted prior to loss of PDMS/collagen adhesion. Samples molded by P-PDMS exhibited increased survival, even when cultured in the presence of TGF-β2. (n = 3, p <0.05; 2-way ANOVA and Tukey’s HSD post hoc test). Mean is indicated by black bars and triangle markers are data points.

### Cultures in porous PDMS molds can be maintained through side wall diffusion

To assess media diffusion in P-PDMS molds, 13 mm tall by 8 mm diameter PDMS molds were used. Both S-PDMS and P-PDMS wells were filled with cell-laden collagen; the P-PDMS molds had interconnected pores(Ren *et al*., 2017; González-Rivera *et al*., 2018; Li *et al*., 2018). Media was added to the top level of the molds but did not cover the top surface, only diffusion through the mold was possible. On day 5, samples were fixed, stained against nuclei, and cleared as described above. Confocal tilescans were acquired of the culture base and stitched into a single volume.

When compared to S-PDMS samples, P-PDMS samples had higher cell densities and well-formed spheroids, indicating robust cell growth (**Figure 6A**). To quantify this, nuclear volume fraction was quantified in both S-PDMS and P-PDMS samples (**Figure 6B & Figure S3**); S-PDMS cultures had a mean nuclear fraction of 0.0045 ± 0.0016, while P-PDMS cultures had a mean value of 0.0210 ± 0.0049 (p = 0.01). To assess media depletion, culture media was isolated at day 1 and day 5 of culture to assay glucose levels (fresh media was added at day 0, day 1, day 2, and day 4). Glucose levels in S-PDMS cultures at days 1 and 5 was 64% and 45% of complete DMEM media at 4.5 g/L, while P-PDMS samples were 16% and 25% of complete media, respectively; these results indicate increased glycolysis of the P-PDMS cultures compared to S-PDMS, consistent with increased cell numbers and metabolism (**Figure 6C**).

**Figure 6:**
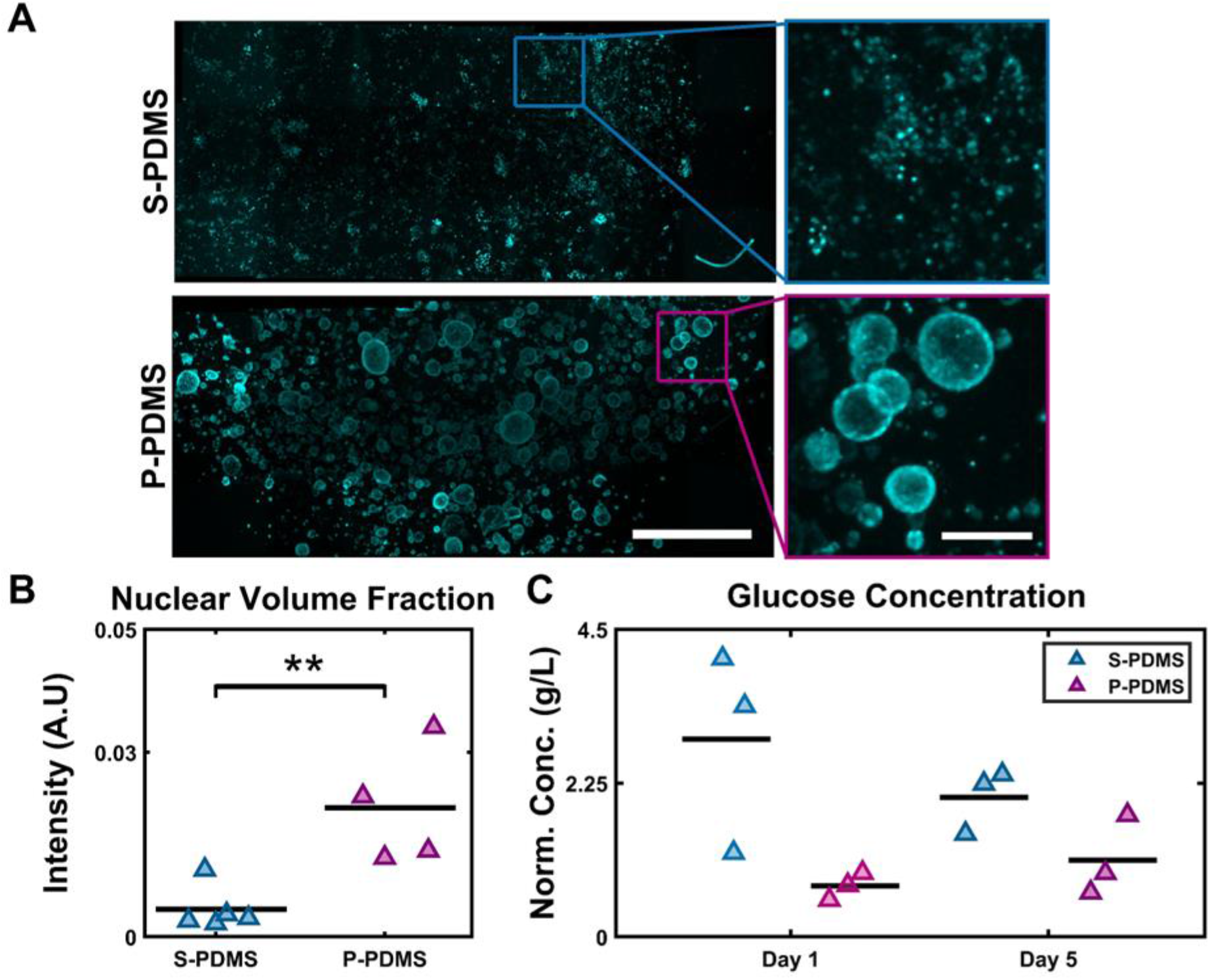
Side feeding through S-PDMS and P-PDMS molds. (A) MDCKs suspended in collagen gels molded by PDA treated S-PDMS or P-PDMS were fed through the side walls. After 7 days of culture, samples were fixed, stained for nuclei and imaged from one end of the culture to the opposite end. Shown are intensity-based maximum projections of stitched tilescans (scalebar = 500 µM) and insets depict zoomed in regions. P-PDMS samples formed distinct spheroids, while S-PDMS sample nuclei remained sparse and scattered. (B) P-PDMS samples had increased nuclear volume fractions relative to S-PDMS (n = 4, p < 0.01; paired sample t-test).(C) Spent culture media was isolated from S-PDMS and P-PDMS samples at days 1 and 5. P-PDMS cultures consumed significantly more glucose than S-PDMS cultures (n = 3, p < 0.05, N-way ANOVA and Tukey’s HSD post hoc test). Mean is indicated by black bars and triangle markers are data points.

#### Porous PDMS anchors VBAM cultures

As an additional demonstration of P-PDMS functionality, we fabricated P-PDMS posts to serve as matrix attachment sites for VBAMs (**Figure 1C**). Briefly, copGFP C2C12s, mCherry HMEC1s, and ASC52telos were suspended in a collagen matrix and seeded around P-PDMS posts inside a S-PDMS chamber at 12.5 million cells/mL total (**Figure 1C**). Despite its importance to tissue maturation, extended culture of dense skeletal muscle constructs remains a challenge due to the contractile nature of the tissue, causing it to rupture and detach from anchor points (Smith *et al*., 2012; Wragg *et al*., 2019; Alave Reyes-Furrer *et al*., 2021). In this study, 24 VBAMs were cultured for 40 days without detachment, prior to downstream processing (**Figure 7A**). For a subset (n = 12) of this population, we observed maturation of the tissues via confocal live imaging of the endogenous fluorophores at days 2 and 21 of differentiation (**Figure 7A**). Early into the differentiation period, muscle cells can be seen stretching out to form myofibers, although void space and rounded cells remain. By day 21 no rounded cells remain, and myofibers are elongated and more compact. We also observed the formation of microvessel-like structures in fixed VBAMs; these structures were predominated localized to the VBAM surface (**Figure 7B-C**).

**Figure 7:**
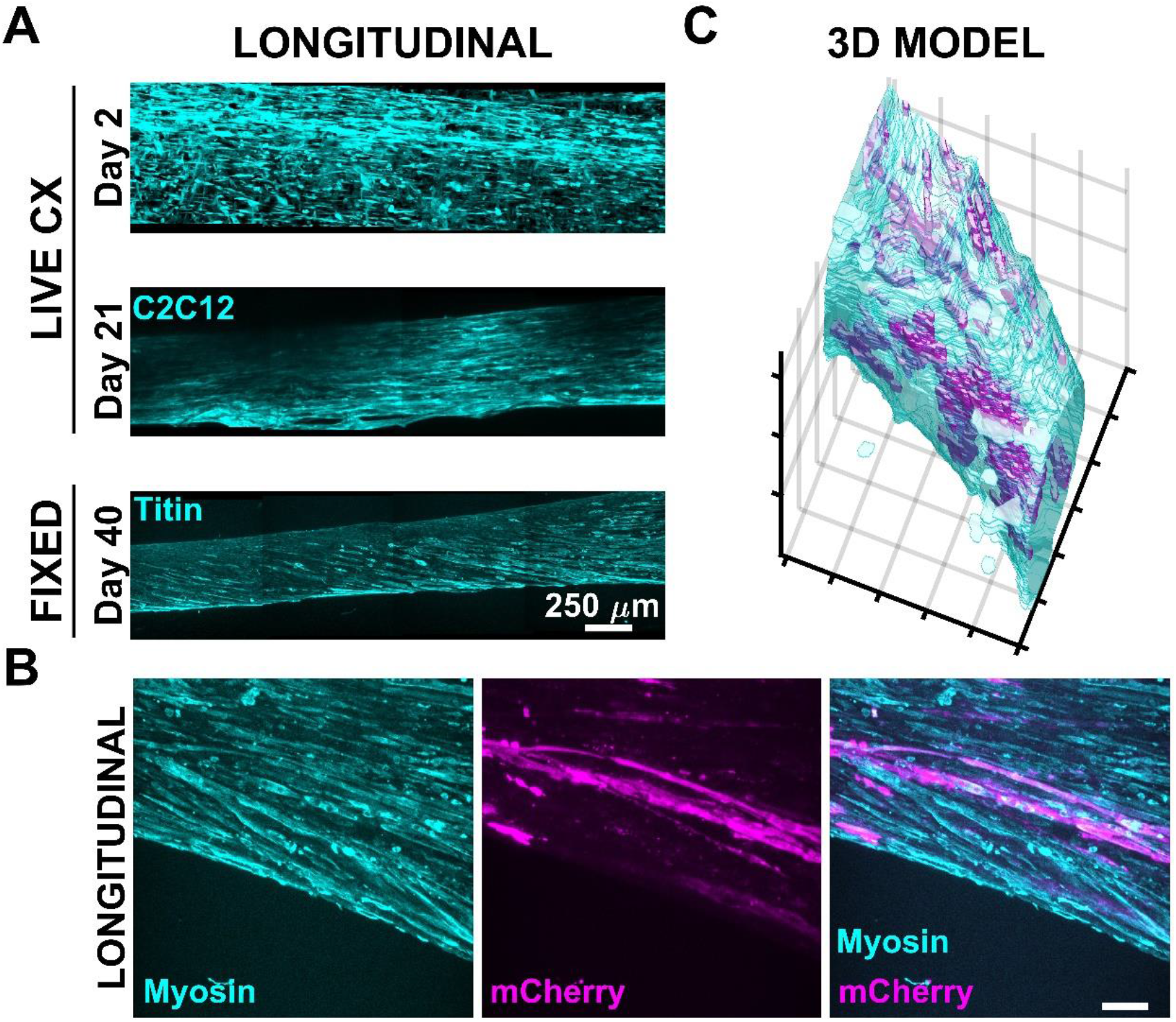
P-PDMS posts support VBAM culture. (A) Shown are intensity-based maximum projections of stitched tilescans encompassing ∼25% of VBAM culture at days 2, 21, and 40 demonstrating stability throughout extended culture. (B) Intensity-based maximum projection of fixed VBAM shows vascular endothelial structures (mCherry) aligning parallel to myofibers (Myosin) (scalebar = 100 µM). (C) Segmented muscle and endothelial channels were used to generate a 3D volume rendering. This 3D model shows the vascular structures are mostly limited to the surface of the VBAM (0.2 mm x 0.4 mm x 0.5mm).

#### Increased feedings do not improve tissue maturation

We additionally tested the impact of feeding frequency on VBAM morphology and maturation. Maturation was evaluated via immunostaining against terminal differentiation markers, myosin (**Figure 8A-B**) or titin (**Figure 8C-D**). In cultures that were fed MWF we observed elongated myofibers with global alignment between the attachment posts (**Figure 8A & 8C**). Additionally, striations are present, consistent with sarcomere formation and functional maturity (Selman Sakar *et al*., 2012; Duffy and Feinberg, 2014; Mueller *et al*., 2021) (**Figure 8A & 8C, insets**). In contrast, B.I.D cultures have myoblasts appeared that have a more rounded shape with little alignment (**Figure 8B & 8D, insets**) and no visible striations; this is characteristic of undifferentiated myoblast cells (**Figure 8B & 8D, insets**).

**Figure 8.**
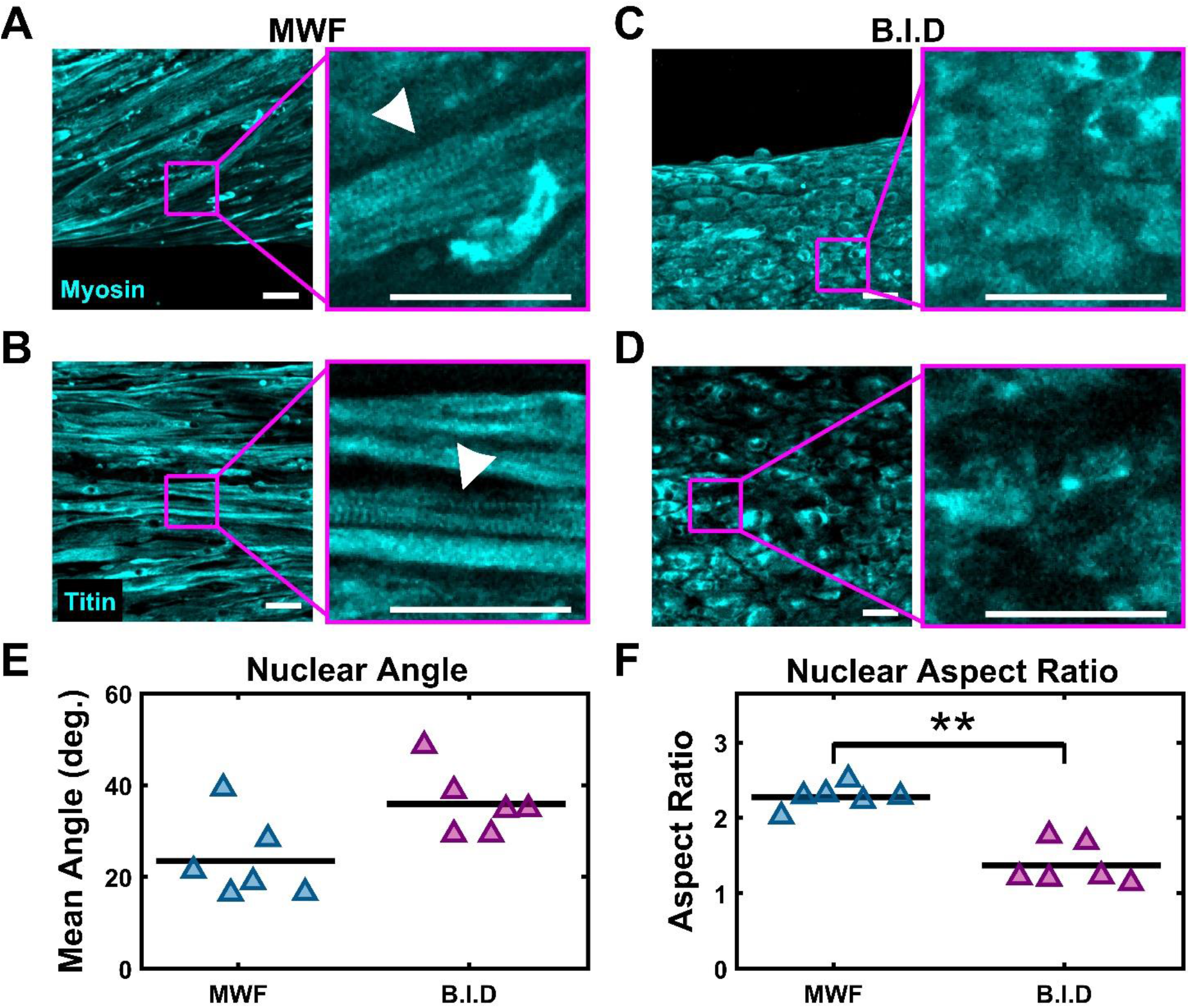
Effects of feeding frequency on VBAM maturation. (A & C) After 5 weeks of culture VBAMs were fixed and stained against myosin (B & D) and titin. Intensity-based maximum projections of MWF VBAMs demonstrate highly aligned myofibers and the presence of striations (insets, white arrowhead). B.I.D VBAMs remained myoblast-like and mononucleated without striations (scalebar = 50 µm). (E) Nuclei in MWF VBAMs have increased alignment relative to B.I.D nuclei (n=6, p = 0.064; paired sample t-test). (F) Nuclei in MWF VBAMs have an increased aspect ratio relative to B.I.D nuclei (n = 6, p < 0.01; paired sample t-test). Mean is indicated by black bars and triangle markers are data points.

To gain insight into VBAM nuclear alignment, we imaged cleared samples and quantified nuclear angle and nuclear aspect ratio. MWF VBAMs have a mean nuclear angle of 23.49° ± 3.64° and B.I.D VBAMs have a mean nuclear angle of 35.93° ± 2.94° (**Figure 8E**). MWF VBAMs have a mean nuclear aspect ratio of 2.27 ± 0.06 and B.I.D VBAMs have a mean value of 1.37 ± 0.11, further indicating MWF VBAMs have improved differentiation (**Figure 8F**).

To qualitatively and quantitatively analyze tissue scale fiber alignment, we analyzed PS-OCT data acquired from the uncleared samples. **Figure 9A-H** shows representative cross-sectional intensity, phase retardation, orientation, and vectorial birefringence images of MWF and B.I.D samples. The intensity images (**Figure 9A & 9E**) show the overall structure of the sample. From the phase retardation images (**Figure 9B & 9F**), we observe that the MWF samples have higher phase retardation (4.28 ± 0.47 rad/mm) than the B.I.D samples (1.78 ± 0.12 rad/mm) which demonstrate a statistically significant difference (p = 0.04) between the two experimental groups (**Figure 9K**). The difference in phase retardation agrees with the difference in morphology of the VBAMs observed in the immunostained images (**Figure 8A-D**). The structural anisotropy arising from the elongated myofibers causes higher phase retardation in the MWF samples. The phase retardation of the MWF samples is higher (>2.8 rad/mm) throughout the length of the samples compared to the B.I.D samples. One interesting characteristic of the MWF samples, is that the phase retardation in the middle of the samples is on average 20% higher compared to the two ends. This suggests that the muscle fibers are more organized and uniformly oriented in the middle as evident from the *en face* phase retardation images (**Figure S2B-C**).

**Figure 9.**
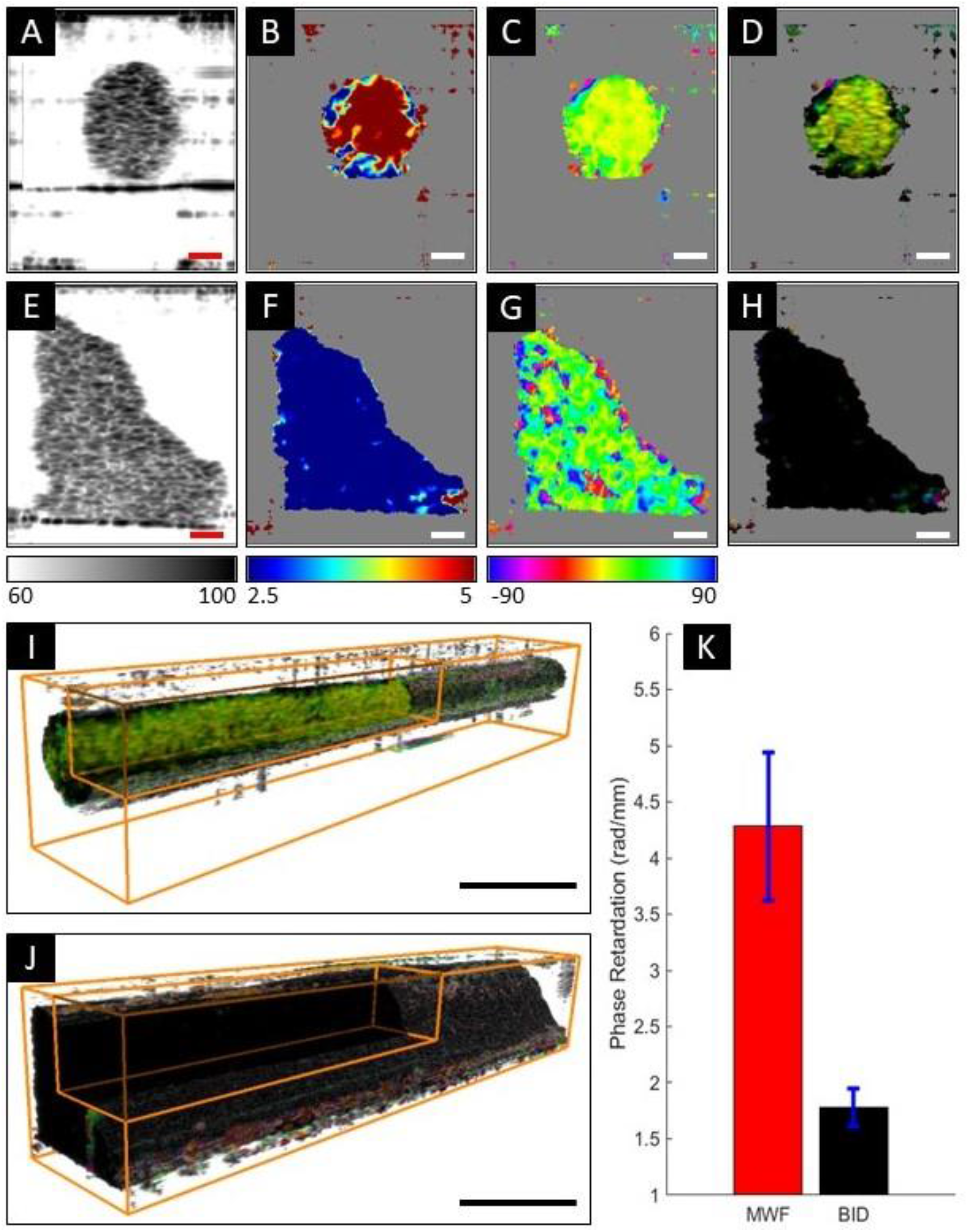
PS-OCT analysis of the effects of feeding frequency on VBAM maturation. (A-D) shows representative cross-sectional intensity (in dB), phase retardation (in rad/mm), orientation (in degree) and vectorial birefringence images, respectively, from MWF sample (scalebar = 100 µm). (E-G) shows the similar cross-sections from B.I.D sample. The MWF samples exhibit higher phase retardation and more uniform orientation compared to the B.I.D samples (scalebar = 100 µm). (I-J) shows representative 3D vectorial birefringence images of 4 mm long segment of MWF and B.I.D samples, respectively (scalebar = 500 µm). The MWF sample has highly organized and uniformly orientated muscle fibers throughout the volume compared to the B.I.D sample. (K) Quantification of the mean phase retardation of the MWF and B.I.D samples statistically improved differentiation (p = 0.04) in the MWF samples.

The orientation images (**Figure 9C & 9G**) also exhibit more uniform fiber orientation in the MWF sample (12.02° ± 14.44°) compared to the B.I.D sample (21.69° ± 35.37°). This is also in line with the immunostained images (**Figure 8A-D**) where we observe well aligned elongated myofibers with striations in the MWF samples compared to rounded myoblasts with little alignment and no striations in the B.I.D samples. The vectorial birefringence images (**Figure 9D & 9H**) combine the phase retardation and the optic axis images using HSV colormap. The cross-sectional vectorial birefringence images were compiled in Amira to generate the 3D vectorial birefringence images (**Figure 9I-J**). The uniformity in color throughout the volume of the MWF sample demonstrates the presence of more highly organized and uniformly orientated muscle fibers than the B.I.D sample.

#### Increased feedings do not improve tissue viability

VBAM cellularity was evaluated using DRAQ7 to label cell nuclei in cleared samples. Volumes were acquired with a confocal microscope and transverse projections of nuclei signal were reconstructed in MATLAB (**Figure 10A**). In MWF tissues cell nuclei are detected throughout the bulk, but there is increased signal at the tissue surface. This indicates that myofibers are preferentially located at the tissue exterior, consistent with cell organization noted in literature (Hinds *et al*., 2011; Ariyasinghe *et al*., 2021). We observed 2 morphologies of nuclear organization in the B.I.D tissues. The first of which is that the constructs remained cellular (**Figure 10A, B.I.D Morph. #1**), but undifferentiated (**Figure 8B** & **8D**). More commonly, there was a considerable decrease in nuclei signal (**Figure 10A, B.I.D Morph. #2**). To better assess the spatial distribution of cells, nuclei signal was segmented, and 3D volume renderings were generated (**Figure 10B & Figure S4**), further confirming that MWF samples had improved cell organization. We additionally tried an intermediate feeding schedule with daily media changes, no improvement in cellularity over MWF was observed (**Figure S5**).

**Figure 10:**
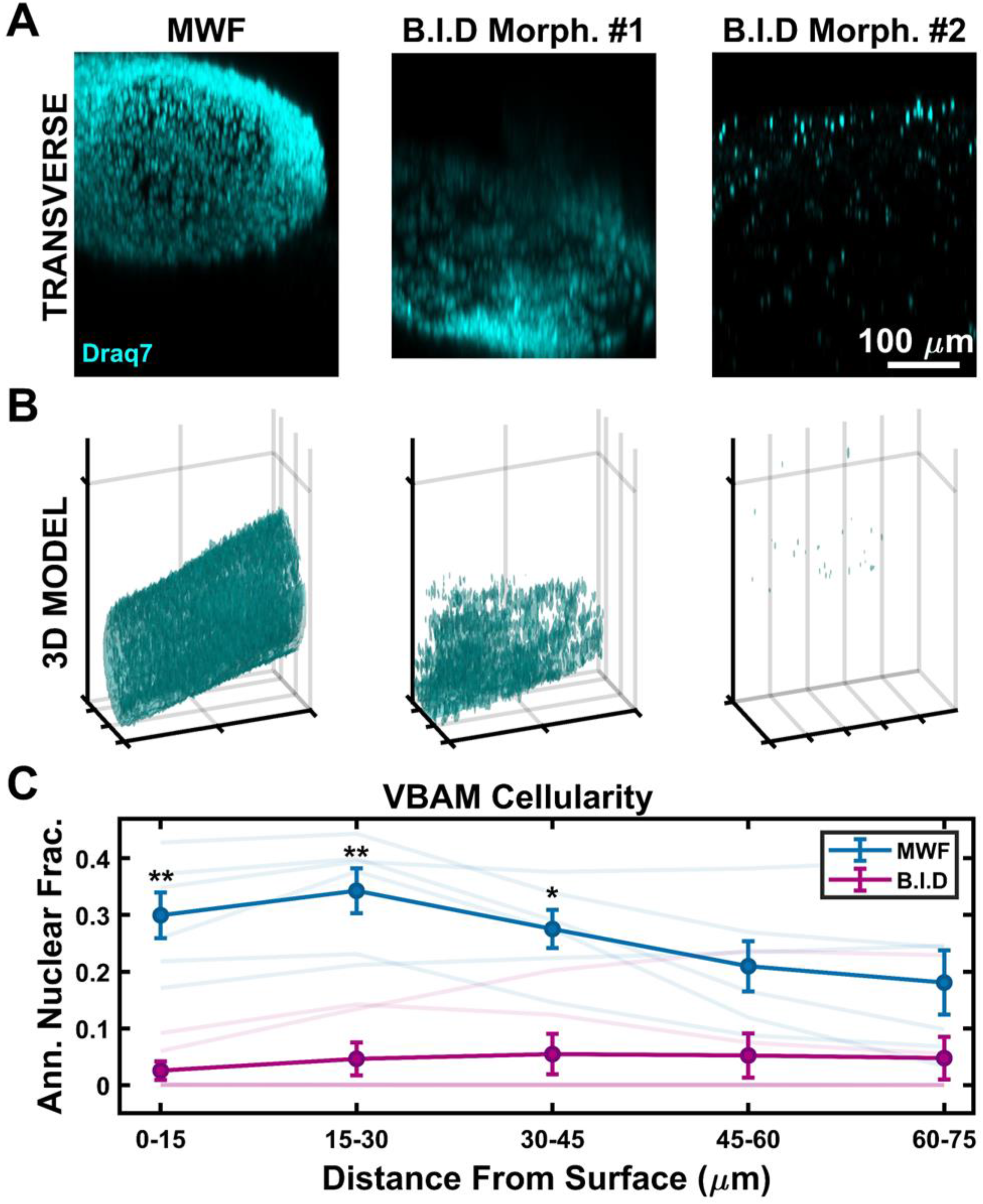
Effects of feeding frequency on VBAM cellularity. (A) Representative transverse projections of VBAM nuclei demonstrate nuclei density and organization throughout the VBAM bulk. MWF VBAMs appear to have increased nuclei density relative to B.I.D morphology #1 and B.I.D morphology #2. (B) 3D volume renderings were generated to visualize the spatial organization of VBAM nuclei, further confirming that MWF VBAMs have increased nuclei density (1 mm x 0.5 mm x 0.6 mm). (C) Annular nuclear fraction was quantified in increments of ∼15 µm radiating inward from the VBAM surface. MWF VBAMs have a significantly higher annular nuclear fraction at 0-15 µm (p < 0.01), 15- 30 µm (p < 0.01), and 30-45 µm (p < 0.05) than B.I.D VBAMs. Mean is indicated by bold blue and magenta lines with +/- SEM shown. Transparent lines show the mean annular nuclear fraction for individual biological replicates (n = 6, paired sample t-test).

To quantify volumetric cellularity, we measured the annular fraction of positively labeled nuclei voxels near the tissue surface (0-15 µm) and deeper into the tissue (15-30 µm, 30-45 µm, 45-60 µm, and 60-75 µm) (**Figure 10C**). In MWF samples, positive annular nuclear fraction at the tissue surface was significantly higher, with a value of 0.30 ± 0.04, relative to B.I.D, with a mean value of 0.03 ± 0.02. This trend continued at 10-20 µm and 20-30 µm deep into the samples, further supporting the observation that more frequent media changes led to decreased tissue cellularity.

## Discussion

Improving 3D cell culture models provides researchers with an important tool for when animal models and traditional 2D culture models are inappropriate. For example, animal models provide important systemic context for disease progression and pharmaceutical response, yet high-throughput and detailed molecular studies can be challenging (Sutherland *et al*., 2013; Brancato *et al*., 2020). Similarly, 2D cell culture models offer increased simplicity and throughput, but lack critical systemic factors and native tissue characteristics (Soares *et al*., 2012; Ravi *et al*., 2015; Costa *et al*., 2016; Jensen and Teng, 2020). Current progress in 3D cell culture models allow for the balancing of increased complexity and physiological relevance with increased control of *in vitro* systems. However, further development of these methods is required to expand applicability to more tissue types and increase researcher access. Key challenges addressed in this study include uncontrolled contraction of the extracellular matrix and mass transfer limitations. Here we introduce a novel culture system for 3D models, capable of supporting highly contractile myofibroblast cells using PDA coated P-PDMS and additionally demonstrate the use of P-PDMS molds to increase cell health in thick cultures. We further apply these techniques to skeletal muscle tissue engineering and demonstrate that PDA coated P-PDMS is a suitable matrix anchor for dense VBAM models. Finally, we evaluate the effect of tissue feeding frequency on overall VBAM health and organization.

Unconstrained cell-mediated gel contraction is a significant challenge with respect to 3D cell culture and tissue engineering (Kosnik *et al*., 2001). Collapse of the extracellular matrix typically occurs in an uncontrolled manner, leading to highly variable outputs for the experiment (Machour *et al*., 2022). In the context of skeletal muscle tissue engineering this problem is particularly detrimental, as it can result in tissue rupture, rendering the construct unusable. While researchers can address this problem by varying the culture length, ECM density, and seeding density, these may reduce maturation, physiological relevance, or otherwise limit the possible experimental conditions (Morgan *et al*., 2019). To address this, 3D culture molds are frequently surface modified to improve ECM adhesion to the mold interface. For instance, PEI/GA surface treatment is often used to improve adhesion of collagen biomaterials to S-PDMS molds (Cross *et al*., 2010), but this linkage still fails when seeding density or matrix stiffness is not optimal (Morgan *et al*., 2019). The bioinspired coating PDA has previously been used to anchor thin biomaterial films to S-PDMS but has not previously been shown to anchor 3D collagen-based cell cultures (Chuah *et al*., 2015; Fu *et al*., 2016; Jeong *et al*., 2016; Lee *et al*., 2018; Harati *et al*., 2022). We evaluated the efficacy of S-PDMS-PEI/GA surface treatments in comparison to S-PDMS-PDA surface treatments and found that PDA coating significantly attenuated cell-mediated gel detachment (**Figure 4**). S-PDMS-PDA is an improved collagen molding method when culturing dense 3D tissue constructs. We further hypothesized that increased surface area of the PDMS molding material would improve collagen adhesion (King *et al*., 2009; Park and Hur, 2021). Prior work has shown P-PDMS can be used as a scaffold for cell culture and cell migration studies (Díaz Lantada *et al*., 2014; Si *et al*., 2016; Zargar *et al*., 2016; Quirós-Solano *et al*., 2018) To demonstrate this in the context of 3D collagen molding, we fabricated P-PDMS and examined PDA coated S-PDMS and P-PDMS culture molds and maintained IMR90 cultures for up to 21 days before terminating the experiment. P-PDMS molding markedly improved collagen attachment (**Figure 5**). P-PDMS as a mold offers improved collagen attachment for longer duration highly contractile cultures.

We were further interested in the impact of P-PDMS on mass transfer. Mass transfer limitations remains a fundamental problem for tissue engineering. Poor diffusion of nutrients and waste products result in cell death deep in cultures and chemotaxis to the more nutrient rich regions of the tissue (Karande *et al*., 2004; Griffith and Swartz, 2006; Buchwald, 2009; Zohar *et al*., 2018; Enrico *et al*., 2022). While bioreactors and tissue perfusion can help resolve this issue, these techniques are not always readily available to non-specialist labs and may not be appropriate for all experiments. Here we describe the use of a P-PDMS mold capable of supporting cultures solely via diffusion through the side wall of the mold, keeping the top of the culture at the air-liquid interface (**Figure 6 & Figure S3**). While still diffusion limited, the easy fabrication of permeable culture molds allows for increasing the scale of a culture or the incorporation of air-liquid interfaces. Potential applications include epithelial and stromal co-cultures where the epithelium is brought to the air-liquid interface, but the culture is too thick to be fed purely through a permeable support (e.g., a cell culture insert).

P-PDMS-PDA is also adaptable for 3D skeletal muscle cultures, a challenging *in vitro* model due to the high contractility. As a proof of concept, we showed that P-PDMS-PDA can be used at matrix attachment points for VBAMs, with no evidence of tissue rupture or detachment after 5 weeks of culture (**Figure 7**). In an effort to improve VBAM maturation and cellularity in the absence of tissue perfusion, we evaluated the effect of media change frequency. We initially hypothesized that more frequent feeding of the VBAM tissues would enhance myotube differentiation (**Figure 8**) and cellular organization (**Figure 10 & Figure S4**), yet we observed an adverse effect. This lack of maturation with increased feeding volume, while surprising, has been observed before (Cooper *et al*., 2004). While the mechanism is unknown, one possibility is the loss of paracrine signals vital to muscle differentiation that accumulate in the culture media; regular removal of those factors too quickly may impair overall VBAM health. Alternatively, the more frequent feeding may mechanically disrupt the cells.

Here, we demonstrated P-PDMS-PDA can support VBAM models with promising cellularity and myotube differentiation, however the vascular cells present in the model failed to form extensive microvessel networks when cultured with muscle cells (**Figure 7**). Instead, microvessel structures were limited to the VBAM surface. This may be due to competition from the densely seeded myoblasts, availability of nutrients, or the increased density of the culture following contraction. Incorporation of perfusion through the VBAM is a promising avenue to resolve nutrient diffusion issues, while providing mechanical signaling that supports microvessel formation (Galie *et al*., 2014; Zohar *et al*., 2018; Kinstlinger *et al*., 2021).

A key advantage of the P-PDMS mold described in this work is the low cost and high accessibility, including the use of commercially available sugar cubes as a sacrificial template. There are drawbacks associated with sugar cube templating, such as fixed pore size based on the diameter of sugar granules, which cannot be readily tuned by the investigator. Should they be required, alternative fabrication strategies are available. P-PDMS can be formed via emulsion templating, phase separation, or use of sacrificial 3D printed templates (Huang *et al*., 2005; Thurgood *et al*., 2017; Montazerian *et al*., 2019). These methods provide more flexibility in mold dimensions and porosity, but generally rely on complex fabrication techniques. Strategies to generate P-PDMS are thoroughly reviewed by Zhu and colleagues (Zhu *et al*., 2017).

Additionally, we provide automated MATLAB algorithms that can be used to volumetrically quantify VBAM nuclei organization, alignment, and morphology in confocal microscopy data (**Supplementary Materials**). Often, these parameters are quantified manually at pre-defined regions within 3D cell constructs, potentially concealing important spatial information. The described automated image analysis methods improve robustness by allowing for the entire 3D bulk to be analyzed and avoiding potential bias that may arise when manually selecting regions to quantify. Measurement of annular nuclear fraction provides further context into the distribution of cells within the tissue. Importantly, these algorithms can be implemented to assess nuclei organization and morphology in a broad range of engineered tissues.

VBAM maturation was further validated with PS-OCT. This imaging modality has previously been used to monitor collagen fiber alignment in engineered tendon constructs (Ahearne *et al*., 2008; Yang *et al*., 2010). In this study, the presence of well aligned elongated myofibers in the MWF samples was evident from the phase retardation and orientation images which conforms with the immunostained images. The 3D vectorial birefringence images, which combine the information of the phase retardation and orientation, demonstrated highly organized and uniformly orientated muscle fibers throughout the volume of the MWF samples compared to the B.I.D samples. The quantitative comparison based on the phase retardation also exhibited improved differentiation in the MWF samples. All these results suggest that PS-OCT is a promising imaging modality for studying engineered tissue because it can non-invasively generate multi-modal 3D volumetric images and allow quantitative analysis without the use of any external agent.

## Conclusion

We have demonstrated a novel culture molding technique capable of supporting contractile and metabolically active engineered tissues by varying S-PDMS surface functionalization and increasing the surface area with P-PDMS molds. P-PDMS molds functionalized with PDA were shown to be suitable anchors for multiple 3D culture types and durations, including VBAMs. Overall, the fabrication methods described are readily extensible to research groups across a broad range of disciplines working on other contractile or metabolically active tissues such as fibrosis models, smooth muscle, and cardiac muscle.

## Supporting information

Supplementary Material; VBAM Annular Nuclear Fraction Code

Supplementary Material; VBAM Nuclear Alignment and Aspect Ratio Code

## Acknowledgements

IAB is supported in part by a TRANSCEND fellowship from the California Institute of Regenerative Medicine (Award # EDUC4-12752). The content is solely the responsibility of the authors and does not necessarily represent the official views of CIRM. This material is partially based upon work supported by the National Science Foundation under Grant No. CAREER 2046093. Any opinions, findings, and conclusions or recommendations expressed in this material are those of the author(s) and do not necessarily reflect the views of the National Science Foundation.

