## Supplementary Material; VBAM Annular Nuclear Fraction Code for "*In vitro* formation and extended culture of highly metabolically active and contractile tissues": bwdistsc.m

```
function D=bwdistsc(bw,aspect)
% D=BWDISTSC(BW,ASPECT)
% BWDISTSC computes Euclidean distance transform of a binary 3D image BW. 
% Distance transform assigns to each pixel in BW a number that is the 
% distance from that pixel to the nearest nonzero pixel in BW. BWDISTSC
% can accept a regular 2D image, a 3D array, and a cell array of 2D slices. 
% ASPECT is a 3-component vector defining the voxel-aspect-ratio for BW. 
% If ASPECT is not given, [1 1 1] isotropic aspect ratio is assumed.
%
% BWDISTSC uses fast optimized scan algorithm and cell-arrays to 
% represent internal data, and is less demanding to physical memory as 
% well as in many cases up to 10 times faster than MATLAB's native bwdist.
%
% Example:
% bw=zeros(100,100,100);
% bw(40:60,40:60,40:60)=1;
% tic;D=bwdist(bw);toc
% tic;D=bwdistsc(bw);toc
%
% BWDISTSC tries to use MATLAB bwdist from image processing toolbox for 2D 
% scans if possible, which is faster, otherwise BWDISTSC will use its own 
% algorithm to also perform 2D scans. Own algorithm is also used if x- and
% y-anisotropy scales are not equal; therefore, if your data has only one
% axis that is anisotropic, it is always advantageous to feed it to
% BWDISTSC so that the anisotropic axis is z.
%
%(c) Yuriy Mishchenko HHMI JFRC Chklovskii Lab JUL 2007
% Updated Yuriy Mishchenko (Toros University) SEP 2013

% This implementation uses optimized forward-backward scan version of the 
% algorithm of the original bwdistsc (2007), which substantially improves
% its speed and simplifies the code. The improvement is described in the 
% part on the selection initial point in the SIVP paper below. The original
% implementation is still used in bwdistsc1, since forward-backward scan
% does not allow limiting computation to a fixed distance value MAXVAL.

% This code is free for use or modifications, just please give credit 
% where appropriate. And if you modify code or fix bugs, please drop 
% me a message at.

%%%%%%%%%%%%%%%%%%%%%%%%%%%%%%%%%%%%%%%%%%%%%%%%%%%%%%%%%%%%%%%%%
% Scan algorithms below use following Lema:                     %
% LEMA: let F(X,z) be lower envelope of a family of parabola:   %
% F(X,z)=min_{i} [G_i(X)+(z-k_i)^2];                            %
% and let H_k(X,z)=A(X)+(z-k)^2 be a parabola.                  %
% Then for H_k(X,z)==F(X,z) at each X there exist at most       %
% two solutions k1<k2 such that H_k12(X,z)=F(X,z), and          %
% H_k(X,z)<F(X,z) is restricted to at most k1<k2.               %
% Here X is any-dimensional coordinate.                         %
%                                                               %
% Thus, simply scan away from any z such that H_k(X,z)<F(X,z)   %
% in either direction as long as H_k(X,z)<F(X,z) and update     %
% F(X,z). Note that need to properly choose starting point;     %
% starting point is any z such that H_k(X,z)<F(X,z); z==k is    %
% usually, but not always the starting point!!!                 %
% usually, but not always the starting point!                   %
%                                                               %
% Citation:                                                     %
% Mishchenko Y. (2013) A function for fastcomputation of large  %
% discrete Euclidean distance transforms in three or more       %
% dimensions in Matlab. Signal, Image and Video Processing      %
% DOI: 10.1007/s11760-012-0419-9.                               %
%%%%%%%%%%%%%%%%%%%%%%%%%%%%%%%%%%%%%%%%%%%%%%%%%%%%%%%%%%%%%%%%%

% parse inputs
if(nargin<2 || isempty(aspect)) aspect=[1 1 1]; end

% determine geometry of the data
if(iscell(bw)) shape=[size(bw{1}),length(bw)]; else shape=size(bw); end

% correct this for 2D data
if(length(shape)==2) shape=[shape,1]; end
if(length(aspect)==2) aspect=[aspect,1]; end
    
% allocate internal memory
D=cell(1,shape(3)); for k=1:shape(3) D{k}=zeros(shape(1:2)); end


%%%%%%%%%%%%% scan along XY %%%%%%%%%%%%%%%%
for k=1:shape(3)    
    if(iscell(bw)) bwXY=bw{k}; else bwXY=bw(:,:,k); end
        
    % initialize arrays
    DXY=zeros(shape(1:2));
    D1=zeros(shape(1:2));

    % if can, use 2D bwdist from image processing toolbox    
    if(exist('bwdist') && aspect(1)==aspect(2))
        D1=aspect(1)^2*bwdist(bwXY).^2;
    else    % if not, use full XY-scan
        %%%%%%%%%%%%%%% X-SCAN %%%%%%%%%%%%%%%        
        % reference for nearest "on"-pixel in bw in x direction down
        
        %  scan bottow-up (for all y), copy x-reference from previous row 
        %  unless there is "on"-pixel in that point in current row, then 
        %  that's the nearest pixel now
        xlower=repmat(Inf,shape(1:2)); 
        
        xlower(1,find(bwXY(1,:)))=1;    % fill in first row
        for i=2:shape(1)
            xlower(i,:)=xlower(i-1,:);  % copy previous row
            xlower(i,find(bwXY(i,:)))=i;% unless there is pixel
        end
        
        % reference for nearest "on"-pixel in bw in x direction up
        xupper=repmat(Inf,shape(1:2));
        
        xupper(end,find(bwXY(end,:)))=shape(1);
        for i=shape(1)-1:-1:1
            xupper(i,:)=xupper(i+1,:);
            xupper(i,find(bwXY(i,:)))=i;
        end
                
        % build (X,Y) for points for which distance needs to be calculated
        idx=find(~bwXY); [x,y]=ind2sub(shape(1:2),idx);
        
        % update distances as shortest to "on" pixels up/down in the above
        DXY(idx)=aspect(1)^2*min((x-xlower(idx)).^2,(x-xupper(idx)).^2);
        
        %%%%%%%%%%%%%%% Y-SCAN %%%%%%%%%%%%%%%
        % this will be the envelop of parabolas at different y
        D1=repmat(Inf,shape(1:2));
        
        p=shape(2);
        for i=1:shape(2)
            % some auxiliary datasets
            d0=DXY(:,i);
            
            % selecting starting point for x:
            % * if parabolas are incremented in increasing order of y, 
            %   then all below-envelop intervals are necessarily right-
            %   open, which means starting point can always be chosen 
            %   at the right end of y-axis
            % * if starting point exists it should be below existing
            %   current envelop at the right end of y-axis
            dtmp=d0+aspect(2)^2*(p-i)^2;
            L=D1(:,p)>dtmp;
            idx=find(L);            
            D1(idx,p)=dtmp(L);
          

            % these will keep track along which X should 
            % keep updating distances            
            map_lower=L;
            idx_lower=idx;
            
            % scan from starting points down in increments of 1
            for ii=p-1:-1:1
                % new values for D
                dtmp=d0(idx_lower)+aspect(2)^2*(ii-i)^2;
                
                % these pixels are to be updated
                L=D1(idx_lower,ii)>dtmp;
                D1(idx_lower(L),ii)=dtmp(L);
                
                % other pixels are removed from scan
                map_lower(idx_lower)=L;                
                idx_lower=idx_lower(L);
                
                if(isempty(idx_lower)) break; end
            end
        end
    end
    D{k}=D1; 
end


%%%%%%%%%%%%% scan along Z %%%%%%%%%%%%%%%%
D1=cell(size(D));
for k=1:shape(3) 
  D1{k}=repmat(Inf,shape(1:2)); 
end

% start building the envelope 
p=shape(3);
for k=1:shape(3)
    % if there are no objects in this slice, nothing to do
    if(isinf(D{k}(1,1)))
      continue;
    end
    
    % selecting starting point for (x,y):
    % * if parabolas are incremented in increasing order of k, then all 
    %   intersections are necessarily at the right end of the envelop, 
    %   and so the starting point can be always chosen as the right end
    %   of the axis
    
    % check which points are valid starting points, & update the envelop
    dtmp=D{k}+aspect(3)^2*(p-k)^2;
    L=D1{p}>dtmp; 
    D1{p}(L)=dtmp(L);    
    
    % map_lower keeps track of which pixels can be yet updated with the 
    % new distance, i.e. all such XY that had been under the envelop for
    % all Deltak up to now, for Deltak<0
    map_lower=L;
        
    % these are maintained to keep fast track of whether map is empty
    idx_lower=find(map_lower);
    
    % scan away from the starting points in increments of -1
    for kk=p-1:-1:1
        % new values for D
        dtmp=D{k}(idx_lower)+aspect(3)^2*(kk-k)^2;
                    
        % these pixels are to be updated
        L=D1{kk}(idx_lower)>dtmp;
        map_lower(idx_lower)=L;
        D1{kk}(idx_lower(L))=dtmp(L);
                    
        % other pixels are removed from scan
        idx_lower=idx_lower(L);
        
        if(isempty(idx_lower)) break; end
    end
end


% prepare the answer
if(iscell(bw))
    D=cell(size(bw));
    for k=1:shape(3) D{k}=sqrt(D1{k}); end
else
    D=zeros(shape);
    for k=1:shape(3) D(:,:,k)=sqrt(D1{k}); end
end

end
```
