## Supplementary Material; VBAM Annular Nuclear Fraction Code for "*In vitro* formation and extended culture of highly metabolically active and contractile tissues": hysteresis3draw.m

```
function [tri,hys]=hysteresis3draw(img,t1,t2,conn)
% function [tri,hys]=HYSTERESIS3D(img,t1,t2,conn)
%
% Hysteresis3d is a simple function that performs trinarisation and
% hysteresis for 2D and 3D images. Hysteresis3d was inspired by Peter
% Kovesi's 2D hysteresis function
% (http://www.csse.uwa.edu.au/~pk/research/matlabfns/). This 3D function
% takes advantage of the 3D connectivities of imfill instead of the 2D
% connectivities of bwselect.
%
% Usage:        [tri,hys]=HYSTERESIS3D(img,t1,t2,conn)
%
% Arguments:    img - image for hysteresis (assumed to be non-negative)
%               t1 - lower threshold value (fraction b/w 0-1, e.g.: 0.1)
%               t2 - upper threshold value (fraction b/w 0-1, e.g.: 0.9)
%                   (t1/t2 can be entered in any order, larger one will be 
%                   set as the upper threshold)
%               conn - number of connectivities (4 or 8 for 2D)
%                                               (6, 18, or 26 for 3D)       
% Returns:
%               tri - the trinarisation image (values are 0, 1, or 2)
%               hys - the hysteresis image (logical mask image)
% 
% Examples:     [tri,hys]=HYSTERESIS3D(img,0.25,0.8,26)
%
% 2012/07/10: written by Luke Xie 
% 2013/12/09: defaults added 
%
% To see an example of hysteresis used to segment a kidney region, please 
% refer to supplement in QSM of Kidney, NMR Biomed, 2013 Dec;26(12):1853-63 
% (http://onlinelibrary.wiley.com/doi/10.1002/nbm.3039/abstract).
% Supplemental material is also available on our CIVMspace: 
% http://www.civm.duhs.duke.edu/lx201204/

%% arguments
if nargin<3
    disp('function needs at least 3 inputs')
    return;
elseif nargin==3
    disp('inputs=3')
    if numel(size(img))==2;
        disp('img=2D')
        disp('conn set at 4 connectivies (number of neighbors)')
        conn=4;
    end
    if numel(size(img))==3; 
        disp('img=3D')
        disp('conn set at 6 connectivies (number of neighbors)')
        conn=6;
    end
end

%% DO NOT scale t1 & t2 based on image intensity range
if t1>t2    % swap values if t1>t2 
	tmp=t1;
	t1=t2; 
	t2=tmp;
end
t1v=t1;
t2v=t2;

%% trinarisation
tri=zeros(size(img));
tri(img>=t1v)=1;
tri(img>=t2v)=2;

%% hysteresis
abovet1=img>t1v;                                     % points above lower threshold
seed_indices=sub2ind(size(abovet1),find(img>t2v));   % indices of points above upper threshold
hys=imfill(~abovet1,seed_indices,conn);              % obtain all connected regions in abovet1 that include points with values above t2
hys=hys & abovet1;
```
