## Supplementary Material; VBAM Annular Nuclear Fraction Code for "*In vitro* formation and extended culture of highly metabolically active and contractile tissues": VBAM_AnnNucFracQuant_dataViz_EXAMPLE_v2023july5.m

```
%%% use with quant script (VBAM_AnnNucFracQuant_EXAMPLE_v2023july5)
%%
%Authors: Isabella A. Bagdasarian, Joshua T. Morgan
%Lab: TIME lab, PI:Dr. Joshua Morgan, Bioengineering Department
%Institution: Univerisity of California, Riverside
%last edited 07/05/2023 IAB

% Description: Reads in pre-processed .mat files containing segmented
% nuclei volumes (Dhs) and muscle bulk distance transform (Kdist). Defines
% annular ring depth and size (m) and quantifies nuclear fraction in each
% ring (ratio). Stores all annular nuclear fraction values for each dataset
% in (nucRatio).

clearvars
close all
clc


%AnnNucFrac Quant data storage location
outdir1 = 'D:\Bella\Muscle\analysis\';
namey = 'AnnNucFrac Quant';
outdir2 = ['output' '_' namey '_' '16-March-2023' '\'];%define saving folder

% load Kdist and segmented nuclei image for annular ring defining
segVars = dir(['D:\Bella\Muscle\analysis\output_AnnNucFrac Quant_08-February-2023\segmentation datasets\', '*.mat']);

PXscaled = 0.3584 * 4; %accounts for 2x impyramid reduction, so should be
%1.4337 for this analysis!


for j = 1:28
    clear Dhs Kdist nucRing m
    BaseFileName = segVars(j).name;
    BaseFileName = BaseFileName(1:end-4);
    
    %load segmented nuclei volume and muscle distance transform
    load(sprintf('%s\\%s\\segmentation datasets\\%s',outdir1,outdir2,segVars(j).name),'Dhs','Kdist');
    
    %upperBound = floor(max(Kdist(:))); identify max distance, useful for new datasets prior to hardcoding
    %number of rings (for example we go 60 pixels deep)
    
    m = [1:10:60]; %hardcoded to go 60 pixels deep with rings 10 pixels wide
     
    clear ratio
    for g = 2:length(m)
        
        % clear ring
        ring = ge(Kdist, m(g-1)) & le(Kdist, m(g));
        trueRing = ring & Dhs;
        
        ratio(g) = sum(trueRing(:))/sum(ring(:));  
        
    end
  
  nucRatio(j,1:6) = ratio; %
    
end


% organize datasets by tissue


%-row 1, 2,3 = 0401_01

%-row 4 = 0401_03

%-row 10 and 11 = 0502_01
%-row 12 to 15 = 0501_03

%-row 18 and 19 = 0506_01

%-row 20 to 22 = 0506_03

% bid
%-row 5 and 6 = 0401_06

%-row 7 to 9 = 0401_07

%-row 16 and 17 = 0502_07

%-row 23-25 = 0506_06

%-row 26 and 27 = 0506_08

%-row 28 = 0502_08

mwfD = [mean([nucRatio(1,:); nucRatio(2,:); nucRatio(3,:)]); ...
    nucRatio(4,:); ...
    mean([nucRatio(10,:); nucRatio(11,:)]);...
    mean([nucRatio(12,:); nucRatio(13,:); nucRatio(14,:); nucRatio(15,:)]);...
    mean([nucRatio(18,:); nucRatio(19,:)]);...
    mean([nucRatio(20,:); nucRatio(21,:); nucRatio(22,:)])];

afD = [mean([nucRatio(5,:); nucRatio(6,:)]); ...
    mean([nucRatio(7,:); nucRatio(8,:); nucRatio(9,:)]); ...
    mean([nucRatio(16,:); nucRatio(17,:)]); ...
    mean([nucRatio(23,:); nucRatio(24,:); nucRatio(25,:)]);
    mean([nucRatio(26,:); nucRatio(27,:)]);...
    nucRatio(28,:)];

% do stats
[h,p] = ttest(mwfD, afD)

% find mean annular intensity for MWF and AF

meanMWF = mean(mwfD);

meanAF = mean(afD);

 % calculate SEM
 
 stdM = std(mwfD);
 stdA = std(afD);
 
 semM = stdM / sqrt(6);
 semA = stdA /sqrt(6);
    
```
