## Supplementary Material; VBAM Annular Nuclear Fraction Code for "*In vitro* formation and extended culture of highly metabolically active and contractile tissues": VBAM_AnnNucFracQuant_EXAMPLE_v2023july5.m

```
%%% use with shell script (VBAM_AnnNucFracQuant_shell_EXAMPLE_v2023july5)
%%
%Authors: Isabella A. Bagdasarian, Joshua T. Morgan
%Lab: TIME lab, PI:Dr. Joshua Morgan, Bioengineering Department
%Institution: Univerisity of California, Riverside
%last edited 07/05/2023 IAB

% Description: Takes in .lif tilescan volumes of engineered VBAM models containing a
% labeled muscle channel and a labeled nuclei channel. Stitches each tilescan 
% into a single volume stored in offsetLoc. Filters and segments muscle bulk
% and saves volumes as .mat files in the target directory fileLoc. Filters
% and segments nuclei and saves volumes as .mat files in the target
% directory fileLoc. Corrects muscle bulk segmention to make in plane with
% nucle channel and saves corrected volume as .mat file in target directory
% fileLoc. Performs distance transform on corrected muscle bulk volume and saves 
% volume as .mat file in target directory fileLoc. Collects an array of
% nuclei morphological parameters including nuclei intensity, area, and
% volume and stores as .mat file in target directory fileLoc.

% Required sub-functions:
% bwdistsc.m
% hysteresis3draw.m


% open file (takes time)
tic;A = bfopen(sprintf('%s%s',dataLoc,IM_FILE));toc;
fprintf('finished opening file \n');
nSeries = length(A);

%% extract metadata
for g = 1:nSeries
    %%
    
    %open OME metadata
    omeMeta = A{1,4}; %the metadata is stored in the 4th column of cells. It is identical for all rows
    series_1 = g;
    series = g - 1;
    
    fprintf(['begin series ' sprintf('%d',series_1) ' analysis...\n'])
    %a complete list of classes that can be pulled is here:
    %https://static.javadoc.io/org.openmicroscopy/ome-xml/6.0.1/ome/xml/meta/MetadataRetrieve.html
    
    %OME metadata is "zero-indexed": the first series is zero
    voxelSizeX = omeMeta.getPixelsPhysicalSizeX(0).value(ome.units.UNITS.MICROMETER).doubleValue();
    voxelSizeY = omeMeta.getPixelsPhysicalSizeY(0).value(ome.units.UNITS.MICROMETER).doubleValue();
    voxelSizeZ = omeMeta.getPixelsPhysicalSizeZ(0).value(ome.units.UNITS.MICROMETER).doubleValue();
    PX1 = omeMeta.getPixelsPhysicalSizeX(0).value().doubleValue(); %physical pixel X dim, um
    PY1 = omeMeta.getPixelsPhysicalSizeY(0).value().doubleValue(); %physical pixel Y dim, um
    PZ = omeMeta.getPixelsPhysicalSizeZ(0).value().doubleValue(); %physical pixel Z dim, um
    SX = omeMeta.getPixelsSizeX(0).getValue(); % image width, pixels
    SY = omeMeta.getPixelsSizeY(0).getValue(); % image height, pixels
    SZ = omeMeta.getPixelsSizeZ(0).getValue(); % number of Z slices
    nP = omeMeta.getPlaneCount(0); %number of planes
    nC = omeMeta.getChannelCount(0); %number of channels
    
    %%
    C1temp = cat(3,A{g,1}{1:nC:nP-2,1}); %channel 1 (myotubes)
    C1{g} = fliplr(C1temp(:,:,:));
    
    
    C2temp = cat(3,A{g,1}{2:nC:nP-1,1}); %channel 2 (mCherry ECs)
    C2{g} = fliplr(C2temp(:,:,:));
    
    C3temp = cat(3,A{g,1}{3:nC:nP,1}); %channel 3 (DRAQ7,nuclei)
    C3{g} = fliplr(C3temp(:,:,:));
    
end

%% STITCH TILESCAN
fprintf('Load offsets & stitched volumes...\n');

if isfile([offsetLoc '\' IM_FILE(1:end-4) '_NucleiOffsets.mat']) && ~REDO_stitch
    load([offsetLoc '\' IM_FILE(1:end-4) '_NucleiOffsets.mat'],'K','R','D','offsetGbl');
    fprintf(1,'Loaded Existing data!\n')
else
    fprintf(1,'Performing Stitching and saving data!\n')
    tic
    %find offsets
    [offsetGbl] = StitchTest_vFeb28th(C3,omeMeta, nSeries);
    
    
    
    %Stitch images
    fprintf(1,'Begin stitching...\n');
    [~, K] = imagestitch_3D(C1, offsetGbl); 
    [~, R] = imagestitch_3D(C2, offsetGbl); 
    [~, D] = imagestitch_3D(C3, offsetGbl); 
    toc
    
    %% reduce volume TWICE for memory management purposes
    
    K = impyramid(impyramid(K,'reduce'),'reduce'); %myosin/titin channel
    R = impyramid(impyramid(R,'reduce'),'reduce'); %mCherry EC channel
    D = impyramid(impyramid(D,'reduce'),'reduce'); %nuclei channel
    
    save([offsetLoc '\' IM_FILE(1:end-4) '_NucleiOffsets.mat'],'K','R','D','offsetGbl');
    
    clearvars A1 A2 bigblend bigstitch Dtemp F M POSI R R2 R3 R4 %clear variables from offset detection script
    
    clearvars C1 C2 C3 C1temp C2temp C3temp %clear temp image vars
end
fprintf(1,'Done Stitching!\n')

%% correct for 4x reduction in volume (via impyramid transform)
PX = PX1 * 4;
PY = PY1 * 4;


%% SEGMENT TISSUE BULK
fprintf('Load segmented muscle volume...\n');

if isfile([fileLoc '\' IM_FILE(1:end-4) '_skmSeg_uncorr.mat']) && ~REDO_skm
    load([fileLoc '\' IM_FILE(1:end-4) '_skmSeg_uncorr.mat'],'Kt','Khs','K1');
    fprintf(1,'Loaded Existing SkMSeg data!\n')
    
else
    fprintf(1,'Begin bulk muscle filtering & segmentation & saving data... \n')
    
    Kmf = medfilt3(K,[3 3 3]);
    
    K1 = imadjustn(Kmf,[0.0; 0.02],[]);
    
    Kth = imtophat(K1,strel('disk',50)); 
    
    
    
    % Hysteresis on filtered volume
    [~,Khs]=hysteresis3draw(Kth,1000,4000,26);
    
    
    
    fprintf(1,'Segmentation complete, cleaning up...\n')
    %
    clearvars Kao
    for  m =  1:length(Khs(1,1,:))
        
        Kao(:,:,m) = bwareaopen(Khs(:,:,m),900); %remove small artifacts
        
    end
    
    Kf = imdilate(Kao,strel('disk',4));
    Kf = imclose(Kf,strel('disk',30));
    Kf = imopen(Kf,strel('disk',10));
    
    clearvars Kt Kf1
    for s = 1:length(Kf(1,1,:))
        
        Kf1(:,:,s) = imfill(Kf(:,:,s),'holes');
        Kt(:,:,s) = bwmorph(Kf1(:,:,s),'thin');
    end
    
    save([fileLoc '\' IM_FILE(1:end-4) '_skmSeg_uncorr.mat'],'Kt','Khs','K1');
    clearvars  Kf1 Kf Kao Kth Kmf
    
end

%% SEGMENT NUCLEI
fprintf('Load segmented nuclei volume...\n');

if isfile([fileLoc '\' IM_FILE(1:end-4) '_nucSeg.mat']) && ~REDO_nuc
    load([fileLoc '\' IM_FILE(1:end-4) '_nucSeg.mat'],'Dhs','D1');
    fprintf(1,'Loaded Existing NucSeg data!\n')
    
else
    fprintf(1,'Begin nuclei filtering & segmentation & saving data... \n')
    
    Dmf = medfilt3(D,[3 3 3]);
    
    Doc = imtophat(Dmf,strel('sphere',12));
    
    D1 = imadjustn(Doc,[0.0000; .02],[]);
    
    D2 = medfilt3(D1,[5 5 5]);
    
    fprintf(1,'Performing 3D nuclei hysteresis segmentation \n')
    
    
    [~,Dhs]=hysteresis3draw(D2,14000,16000,26);
    
    Dhs=bwareaopen(Dhs,6,26);
    
    save([fileLoc '\' IM_FILE(1:end-4) '_nucSeg.mat'],'Dhs','D1');
    
    clearvars Dmf D2 Doc D1 
    
end

%% CORRECT MUSCLE SEGMENTATION
fprintf('Load corrected muscle segmentation volume...\n');

if isfile([fileLoc '\' IM_FILE(1:end-4) '_skmSeg_corr.mat']) && ~REDO_skmCorr
    load([fileLoc '\' IM_FILE(1:end-4) '_skmSeg_corr.mat'],'collE');
    fprintf(1,'Loaded Existing SkmSeg_corr data!\n')
    
else
    fprintf(1,'Begin correcting muscle segmentation & saving data... \n')
    j = 1;
    
    collE = Kt;
    
    while le(j,numErode(1))
        
        nErode = imerode(collE, strel('sphere', 1));
        
        %counter erode- define a shell of what was just eroded
        shellE = ~nErode & collE;
        
        %check inside shell for nuclei
        testN = shellE & Dhs;
        
        ratio(j) = sum(testN(:))/sum(shellE(:));
        % disp(ratio);
        
        collE = nErode; %update erosion variable
        
        j = j+1;
    end
    
    save([fileLoc '\' IM_FILE(1:end-4) '_skmSeg_corr.mat'],'collE');
    
end

%% PERFORM DISTANCE TRANSFORM
fprintf('Load corrected muscle segmentation volume...\n');

if isfile([fileLoc '\' IM_FILE(1:end-4) '_distTrans.mat']) && ~REDO_dist
    load([fileLoc '\' IM_FILE(1:end-4) '_distTrans.mat'],'Kdist');
    fprintf(1,'Loaded Existing distance transform data!\n')
    
else
    fprintf(1,'Begin performing distance transform & saving data... \n')
    
    Km = ~collE;
    
    Kdist = bwdistsc(Km,[1 1 PZ/PX]);
    
    save([fileLoc '\' IM_FILE(1:end-4) '_distTrans.mat'],'Kdist');
end

%% collect regionprops3 data on nuclei channel

CC = bwconncomp(Dhs,26); 

fprintf(1,'Begin 2D data collection...\n');

NucStats=regionprops3(CC,Dhs,'Volume','Centroid','VoxelList','EquivDiameter','PrincipalAxisLength'); 
NucStats = table2struct(NucStats);

for m = 1:size(Dhs,3)
    stats2D = regionprops(Dhs(:,:,m),D(:,:,m),'MeanIntensity','Area');
    int2D{m} = [stats2D.MeanIntensity]';
    area2D{m} = [stats2D.Area]';
    
    integInt{m} = int2D{m} .* area2D{m};
end

label=labelmatrix(CC);
countNuc=CC.NumObjects;

save([fileLoc '\' IM_FILE(1:end-4) '_DATA.mat'],'int2D','area2D','countNuc','integInt');

%% aggregate major segmentation variables and save- for easier downstream processing
distScale = PZ/PX;

save([fileLoc '\' IM_FILE(1:end-4) '_segVars.mat'],'collE','Khs','Dhs','Kdist','PX','distScale','-v7.3');

%% render segmentation visualization
if REDO_segVis
    
    fprintf('Rendering filtered nuclei channel...\n');
    
    figure
    set(gcf,'units','inches','position',[1 1 5 5],'color','white')
    for i = 1:SZ
        IM = cat(3, zeros(size(D1(:,:,i))),D1(:,:,i),D1(:,:,i));
        image(IM); axis off equal; set(gca,'position',[0 0 1 1]); hold on
        
        %Convert the figure window as displayed into a "frame"
        fr(i) = getframe(gcf);
        
        % To generate a GIF use:
        %     to ensure consistent coloration when we create the GIF (this is
        %     specific to indexed formats like GIF, we create a single colormap
        if i == 1
            [im,map] = rgb2ind(fr(i).cdata,colorcube(256),'nodither'); %define a colormap to be used for the GIF
            im(1,1,1,SZ) = 0; %this just preallocates a matrix of appropriate size for the rest of the loop
        end
        
        %assign indexed data to an image plane
        im(:,:,1,i) = rgb2ind(fr(i).cdata,map,'nodither');
        
        
    end
    close(gcf)
  
    imwrite(im,map,(sprintf('%s\\%s_draqChannel.gif',fileLoc, BaseFileName)),'DelayTime',0.08,'LoopCount',inf)
    
    fprintf('Rendering segmented nuclei channel...\n');
    
    figure
    set(gcf,'units','inches','position',[1 1 5 5],'color','white')
    for i = 1:SZ
        
        imagesc(Dhs(:,:,i)); axis off equal; set(gca,'position',[0 0 1 1]); hold on
        
        %Convert the figure window as displayed into a "frame"
        fr(i) = getframe(gcf);
        
        % To generate a GIF use:
        %     to ensure consistent coloration when we create the GIF (this is
        %     specific to indexed formats like GIF, we create a single colormap
        if i == 1
            [im,map] = rgb2ind(fr(i).cdata,colorcube(256),'nodither'); %define a colormap to be used for the GIF
            im(1,1,1,SZ) = 0; %this just preallocates a matrix of appropriate size for the rest of the loop
        end
        
        %assign indexed data to an image plane
        im(:,:,1,i) = rgb2ind(fr(i).cdata,map,'nodither');
        
        
    end
    close(gcf)
    
    imwrite(im,map,(sprintf('%s\\%s_draqChannelSEG.gif',fileLoc,BaseFileName)),'DelayTime',0.08,'LoopCount',inf)
    
    
    fprintf('Rendering filtered muscle channel...\n');
    
    figure
    set(gcf,'units','inches','position',[1 1 5 5],'color','white')
    for i = 1:SZ
        IM = cat(3, zeros(size(K1(:,:,i))),K1(:,:,i),K1(:,:,i));
        image(IM); axis off equal; set(gca,'position',[0 0 1 1]); hold on
        
        %Convert the figure window as displayed into a "frame"
        fr(i) = getframe(gcf);
        
        % To generate a GIF use:
        %     to ensure consistent coloration when we create the GIF (this is
        %     specific to indexed formats like GIF, we create a single colormap
        if i == 1
            [im,map] = rgb2ind(fr(i).cdata,colorcube(256),'nodither'); %define a colormap to be used for the GIF
            im(1,1,1,SZ) = 0; %this just preallocates a matrix of appropriate size for the rest of the loop
        end
        
        %assign indexed data to an image plane
        im(:,:,1,i) = rgb2ind(fr(i).cdata,map,'nodither');
        
        
    end
    close(gcf)
   
    imwrite(im,map,(sprintf('%s\\%s_SkMChannel.gif',fileLoc, BaseFileName)),'DelayTime',0.08,'LoopCount',inf)
    
    fprintf('Rendering segmented muscle channel...\n');
    
    figure
    set(gcf,'units','inches','position',[1 1 5 5],'color','white')
    for i = 1:SZ
        
        image(collE(:,:,i)); axis off equal; set(gca,'position',[0 0 1 1]); hold on
        
        %Convert the figure window as displayed into a "frame"
        fr(i) = getframe(gcf);
        
        % To generate a GIF use:
        %     to ensure consistent coloration when we create the GIF (this is
        %     specific to indexed formats like GIF, we create a single colormap
        if i == 1
            [im,map] = rgb2ind(fr(i).cdata,colorcube(256),'nodither'); %define a colormap to be used for the GIF
            im(1,1,1,SZ) = 0; %this just preallocates a matrix of appropriate size for the rest of the loop
        end
        
        %assign indexed data to an image plane
        im(:,:,1,i) = rgb2ind(fr(i).cdata,map,'nodither');
        
        
    end
    close(gcf)

    imwrite(im,map,(sprintf('%s\\%s_SkMChannelSEG.gif',fileLoc,BaseFileName)),'DelayTime',0.08,'LoopCount',inf)
    
    fprintf('Rendering segmentation tracings...\n');
    
    figure
    set(gcf,'units','inches','position',[1 1 6 6],'color','white')
    for i = 1:SZ
        IM = cat(3, zeros(size(D1(:,:,i))),K1(:,:,i),D1(:,:,i));
         %CREATE AN AXIS THAT FILLS THE ENTIRE FIGURE
    
        %display the image plane and clear off the axis and tick marks
        image(IM); set(gca,'visible','off'); axis equal; hold on
        
       %  image(IM); axis off equal; set(gca,'position',[0 0 1 1]); hold on
        BG1 = bwboundaries(Dhs(:,:,i));
        for f = 1:length(BG1)
            plot(BG1{f}(:,2),BG1{f}(:,1),'w');
        end
          
        BG3 = bwboundaries(collE(:,:,i));
        for f = 1:length(BG3)
            plot(BG3{f}(:,2),BG3{f}(:,1),'color','m');
        end
        
        %Convert the figure window as displayed into a "frame"
        fr(i) = getframe(gcf);
        
        
        % To generate a GIF use:
        %     to ensure consistent coloration when we create the GIF (this is
        %     specific to indexed formats like GIF, we create a single colormap
        if i == 1
            [im,map] = rgb2ind(fr(i).cdata,colorcube(256),'nodither'); %define a colormap to be used for the GIF
            im(1,1,1,SZ) = 0; %this just preallocates a matrix of appropriate size for the rest of the loop
        end
        
        %assign indexed data to an image plane
        im(:,:,1,i) = rgb2ind(fr(i).cdata,map,'nodither');
        
    
    end
    close(gcf)
 
    imwrite(im,map,(sprintf('%s\\%s_segTracings.gif',fileLoc, BaseFileName)),'DelayTime',0.08,'LoopCount',inf)
    
    
end

if REDO_3D
    
    [Giso] = myoVolRendering_v2021oct19(dataLoc,IM_FILE,Dhs,outdir1,...
        outdir2,PX,PY);
    
    save(sprintf('%s\\%s_3DVolRender.mat',fileLoc,BaseFileName),'Giso','distScale','-v7.3');
    
end
```
