## Supplementary Material; VBAM Nuclear Alignment and Aspect Ratio Code for "*In vitro* formation and extended culture of highly metabolically active and contractile tissues": VBAM_NucAlignment_dataViz_EXAMPLE_v2023july5.m

```
%%% use with quant script (VBAM_NucAlignmentQuant_EXAMPLE_v2023july5)
%%
%Authors: Isabella A. Bagdasarian, Joshua T. Morgan
%Lab: TIME lab, PI:Dr. Joshua Morgan, Bioengineering Department
%Institution: Univerisity of California, Riverside
%last edited 07/05/2023 IAB

% Description: Reads in pre-processed .mat files containing nuclei angles (NDth). 
% Stores all nuclei angle values for each dataset in (normDirAll).

clearvars
close all
clc

%load output NucStats data from VBAM_NucAlignmentQuant
dataVars = dir(['D:\Bella\Muscle\analysis\output_NucAlignment Quant_20-February-2023\data outputs\', '*.mat']);

% FIND NUCLEI ASPECT RATIO
for j = 1:length(dataVars)
    
        
        clear NDth;
        BaseFileName = dataVars(j).name;
        BaseFileName = BaseFileName(1:end-4);
        load(sprintf('%s\\data outputs\\%s','D:\Bella\Muscle\analysis\output_NucAlignment Quant_30-April-2023',dataVars(j).name),'NDth');
        
        normDirAll{j} = NDth'; 
        
end
```
