## Supplementary Material; VBAM Nuclear Alignment and Aspect Ratio Code for "*In vitro* formation and extended culture of highly metabolically active and contractile tissues": VBAM_NucAlignmentQuant_EXAMPLE_v2023july5.m

```
%%% use with shell script (VBAM_NucAlignmentQuant_shell_EXAMPLE_v2023july5)
%%
%Authors: Isabella A. Bagdasarian, Joshua T. Morgan
%Lab: TIME lab, PI:Dr. Joshua Morgan, Bioengineering Department
%Institution: Univerisity of California, Riverside
%last edited 07/05/2023 IAB

% Description: Takes in .lif tilescan volumes of engineered VBAM models containing a
% labeled muscle channel and a labeled nuclei channel. Stitches each tilescan 
% into a single volume stored in offsetLoc. Filters and segments muscle bulk
% and saves volumes as .mat files in the target directory fileLoc. Filters,
% segments, and watersheds nuclei and saves volumes as .mat files in the target
% directory fileLoc.  Collects an eigenvectors and eigen values of segmented 
% nuclei and muscle volumes and stores .mat files in the target directory fileLoc
% Performs skeletonization on segmented muscle volume to identify bulk
% tissue direction and saves as .mat files in the target directory fileLoc.
% Calculates local direction of nuclei eigenvectors relative to muscle
% skeleton and corrects for parallel and anti-parallel vectors (NDth) saves as
% .mat files in the target directory fileLoc.

% Required sub-functions:
% imthbr.m
% hysteresis3draw.m


% open file (takes time)
tic;A = bfopen(sprintf('%s%s',dataLoc,IM_FILE));toc;
fprintf('finished opening file \n');
nSeries = length(A);

%% extract metadata
for g = 1:nSeries
    %%
    
    %open OME metadata
    omeMeta = A{1,4}; %the metadata is stored in the 4th column of cells. It is identical for all rows
    series_1 = g;
    series = g - 1;
    
    fprintf(['begin series ' sprintf('%d',series_1) ' analysis...\n'])
    %a complete list of classes that can be pulled is here:
    %https://static.javadoc.io/org.openmicroscopy/ome-xml/6.0.1/ome/xml/meta/MetadataRetrieve.html
    
    %OME metadata is "zero-indexed": the first series is zero
    %get the pixel size in um:
    voxelSizeX = omeMeta.getPixelsPhysicalSizeX(0).value(ome.units.UNITS.MICROMETER).doubleValue();
    voxelSizeY = omeMeta.getPixelsPhysicalSizeY(0).value(ome.units.UNITS.MICROMETER).doubleValue();
    voxelSizeZ = omeMeta.getPixelsPhysicalSizeZ(0).value(ome.units.UNITS.MICROMETER).doubleValue();
    PX1 = omeMeta.getPixelsPhysicalSizeX(0).value().doubleValue(); %physical pixel X dim, um
    PY = omeMeta.getPixelsPhysicalSizeY(0).value().doubleValue(); %physical pixel Y dim, um
    PZ = omeMeta.getPixelsPhysicalSizeZ(0).value().doubleValue(); %physical pixel Z dim, um
    SX = omeMeta.getPixelsSizeX(0).getValue(); % image width, pixels
    SY = omeMeta.getPixelsSizeY(0).getValue(); % image height, pixels
    SZ = omeMeta.getPixelsSizeZ(0).getValue(); % number of Z slices
    nP = omeMeta.getPlaneCount(0); %number of planes
    nC = omeMeta.getChannelCount(0); %number of channels
    
    %%
    C1temp = cat(3,A{g,1}{1:nC:nP-2,1}); %channel 1 (myotubes)
    C1{g} = fliplr(C1temp(:,:,:));
    
    
    C2temp = cat(3,A{g,1}{2:nC:nP-1,1}); %channel 2 (mCherry ECs)
    C2{g} = fliplr(C2temp(:,:,:));
    
    C3temp = cat(3,A{g,1}{3:nC:nP,1}); %channel 3 (DRAQ7,nuclei)
    C3{g} = fliplr(C3temp(:,:,:));
    
end

%% STITCH TILESCAN
fprintf('Load offsets & stitched volumes...\n');

if isfile([offsetLoc '\' IM_FILE(1:end-4) '_NucleiOffsets.mat']) && ~REDO_stitch
    imOffsets = load([offsetLoc '\' IM_FILE(1:end-4) '_NucleiOffsets.mat'],'offsetGbl'); %#ok<*NASGU>
    fprintf(1,'Loaded Existing data!\n')
    offsets = {};
    for f = 1:length(imOffsets)
        offsets{f} = (imOffsets(f).offsetGbl); %remove offsets from structure into cell array
    end
    offsetGbl = vertcat(offsets{:}); %remove offsets from cell array into nx3 doube array
    
    %% Stitch images
    fprintf(1,'Begin stitching...\n'); tic
    [~, K] = imagestitch_3D(C1, offsetGbl); toc 
    [~, R] = imagestitch_3D(C2, offsetGbl); toc 
    [~, D] = imagestitch_3D(C3, offsetGbl); toc 
    
    
    clearvars C1 C2 C3 C1temp C2temp C3temp
    fprintf(1,'Stitching complete!\n');
    
else
    fprintf(1,'Performing Stitching and saving data!\n')
    tic
    %find offsets
    [offsetGbl] = StitchTest_vFeb28th(C3,omeMeta, nSeries);
    
    
    
    %Stitch images
    fprintf(1,'Begin stitching...\n');
    [~, K] = imagestitch_3D(C1, offsetGbl); 
    [~, R] = imagestitch_3D(C2, offsetGbl); 
    [~, D] = imagestitch_3D(C3, offsetGbl); 
    toc
    
    save([offsetLoc '\' IM_FILE(1:end-4) '_NucleiOffsets.mat'],'K','R','D','offsetGbl','-v7.3');
    
    clearvars A1 A2 bigblend bigstitch Dtemp F M POSI R R2 R3 R4 %clear variables from offset detection script
    
    clearvars C1 C2 C3 C1temp C2temp C3temp %clear temporary image storage variables
end
fprintf(1,'Done Stitching!\n')


%% SEGMENT NUCLEI
fprintf('Load segmented nuclei volume...\n');

if isfile([fileLoc '\' IM_FILE(1:end-4) '_nucSeg.mat']) && ~REDO_nuc
    load([fileLoc '\' IM_FILE(1:end-4) '_nucSeg.mat'],'Dhs','D1','Dhs_10','Dhs_12','D2','Doc');
    fprintf(1,'Loaded Existing NucSeg data!\n')
    
else
    fprintf(1,'Begin nuclei filtering & segmentation & saving data... \n')
    
    Dmf = medfilt3(D,[3 3 3]);
    
    Doc = zeros(size(Dmf),'uint16'); 
    for i = 1:SZ
        Doc (:,:,i) = imthbr(Dmf(:,:,i),strel('disk',10));
    end
    
    D1 = imadjustn(Doc,[.0000; .02],[]);
    D2 = medfilt3(D1,[5 5 5]);
    
    %for initial segmentation collect single nuclei, rather than collecting all nuclei
    
    [~,Dhs]=hysteresis3draw(D2,16000,20000,26);
       
    save([fileLoc '\' IM_FILE(1:end-4) '_nucSeg.mat'],'Dhs','D1','D2','Doc','-v7.3');
    
    clearvars Dmf D2
    
end

%% WATERSHED NUCLEI
fprintf('Load watershedded nuclei volume...\n');

if isfile([fileLoc '\' IM_FILE(1:end-4) '_nucWS.mat']) && ~REDO_WS
    load([fileLoc '\' IM_FILE(1:end-4) '_nucWS.mat'],'Dlg2');
    fprintf(1,'Loaded Existing NucWS data!\n')
    
else
    fprintf(1,'Begin nuclei watershedding... \n')
    
    % Watershed Nuclei and remove non-segmented objects
    tic;
    ED = -bwdist(~Dhs); toc %create a euler distance map (the distance from any pixel to the nearest background)
    %the minus sign converts each nuclear "hill" (since the center is furthest from the edge) to a "valley"
    ED(~Dhs) = Inf;
    
    ED = imhmin(ED,1); %sometimes you can have 2 barely different minimas, the hmin will suppress any small minima
    tic
    Dws = watershed(ED); toc %watershedding associates each point with its nearest minima. Think of it like a drop of water hitting each pixel, which drops flow together?
    Dws(~Dhs) = 0;
    
    % zero out non segmented objects
    Dws(Dhs == 0) = 0;
    
    % convert to a logical
    Dlg = logical(Dws);
    
    %smooth objects
    Dlg2 = imopen(Dlg,strel("disk",3));
    
    save([fileLoc '\' IM_FILE(1:end-4) '_nucWS.mat'],'Dlg2','-v7.3');
    
    clear D D1 Dhs Dlg Doc Dws
    
end

%% PERFORM MUSCLE SEGMENTATION
fprintf('Load muscle segmentation volume...\n');

if isfile([fileLoc '\' IM_FILE(1:end-4) '_skmSeg_corr.mat']) && ~REDO_skm
    load([fileLoc '\' IM_FILE(1:end-4) '_skmSeg_corr.mat'],'Khs','Kmf','K1','Koc2');
    fprintf(1,'Loaded Existing SkmSeg data!\n')
    
else
    fprintf(1,'Begin 3D muscle filter segmentation & saving data... \n')
    
    tic
    Kmf = medfilt3(K,[3 3 3]); toc
    
    tic
    K1 = imadjustn(Kmf,[0.0; 0.02],[]); toc
    
    Koc = zeros(size(K1),'uint16');
    for i = 1:SZ
        Koc (:,:,i) = imthbr(K1(:,:,i),strel('disk',500));
    end
    
    Koc2 = imclose(Koc, strel('disk',5));
    
    [~,Khs]=hysteresis3draw(Koc2,5000,10000,26);
    
    Khs = bwareaopen(Khs,1e7);
    
    save([fileLoc '\' IM_FILE(1:end-4) '_skmSeg_corr.mat'],'Khs','Kmf','K1','Koc2','-v7.3');
    clear K1 Kmf Koc Koc2 K
    
end

%% collect regionprops3 data
fprintf('finding nuclei and muscle eigenvectors...\n');
L = bwlabeln(Dlg2);
NucStats=regionprops3(L,'EigenVectors','EigenValues','Orientation','PrincipalAxisLength','Centroid');
cx = NucStats.Centroid(:,1); cy = NucStats.Centroid(:,2); cz = NucStats.Centroid(:,3);
NucStats = table2struct(NucStats);
NucDir = zeros(length(NucStats),3);
NumNuc = length(NucStats);
for i = 1:length(NucStats)
    NucDir(i,1:3) = NucStats(i).EigenVectors(:,1)';
end
U = NucDir(:,1);
V = NucDir(:,2);
W = NucDir(:,3);
X = zeros(length(NucStats),1);

clear A L ED R

s = regionprops3(Khs,'EigenVectors','EigenValues','Orientation','PrincipalAxisLength');
MuscDirVolume = s.EigenVectors{1}(:,1)';

fprintf('Load muscle skeleton volume & angles...\n');


try 
    load([fileLoc '\' IM_FILE(1:end-4) '_skmSKEL.mat'],'S2','GD1','GD2','GDm','Cost');
    fprintf(1,'Loaded Existing SkmSKEL data!\n')
    
catch
    %complex muscdir by finding centerline
    %first, clean up the object
    fprintf('Filter segmented muscle volume...\n');
    Khs2 = imclose(Khs,strel('disk',20));
    Khs2 = imdilate(Khs2,strel('disk',10)); % clean edges
    Khs2 = imopen(Khs2,strel('disk',50));
        
    %perform bwdist but completely ignore 3rd dimension, too much noise from
    %the lumpy upper/lower surface of the segmentation
    fprintf('Perform distance transform...\n');
    tic
    Dst = zeros(size(Khs2));
    c1out = zeros(size(Khs2,3),2);
    c2out = zeros(size(Khs2,3),2);
    fd = zeros(size(Khs2,3),1);
    for i = 1:size(Khs2,3)
        Dst(:,:,i)=bwdist(~Khs2(:,:,i));
        st = bwferet(Khs2(:,:,i),"MaxFeretProperties");
        if ~isempty(st)
            %Feret reports as 0.5 sometimes, so round
            c1 = round(st.MaxCoordinates{1}(1,:));
            c2 = round(st.MaxCoordinates{1}(2,:));
            %force over dimension coordinates on map
            c1 = min(c1,[size(Khs2,2) size(Khs2,1)]);
            c2 = min(c2,[size(Khs2,2) size(Khs2,1)]);
            
            c1out(i,:) = c1; c2out(i,:) = c2;
            fd(i) = max(st.MaxDiameter);
        else
            fd(i) = -Inf;
        end
    end
    toc
    
    %the above checks each plane, but maybe it makes more sense to flatten the
    %image, just in case the full span isn't in a single plane
    st = bwferet(max(Khs2,[],3),"MaxFeretProperties");
    %Feret reports as 0.5 sometimes, so round
    c1 = round(st.MaxCoordinates{1}(1,:));
    c2 = round(st.MaxCoordinates{1}(2,:));
    %force over dimension coordinates on map
    c1 = min(c1,[size(Khs2,2) size(Khs2,1)]);
    c2 = min(c2,[size(Khs2,2) size(Khs2,1)]);
    
    %create SEED arrays that are true around dilations of the feret points
    SEED1 = false(size(Khs2,1),size(Khs2,2)); SEED1(c1(2),c1(1)) = true;
    SEED1 = imdilate(SEED1,strel('disk',50,0));
    
    SEED2 = false(size(Khs2,1),size(Khs2,2)); SEED2(c2(2),c2(1)) = true;
    SEED2 = imdilate(SEED2,strel('disk',50,0));
    
    %scale SEED arrays
    SEED1 = repmat(SEED1,[1 1 size(Khs2,3)]);
    SEED2 = repmat(SEED2,[1 1 size(Khs2,3)]);
    
    LTEMP = bwlabeln(Khs2); 
    
    Rprops = regionprops3(LTEMP,'Volume'); 
    SEED2d = imdilate(SEED2,strel('disk',70,0));     
    temp = unique(LTEMP(SEED2d));
    
    clear Khs Khs2 ED

    fprintf('Find cost matrix...\n');
    %create the cost matrix from Dst, strongly favor centerline using -4 power
    maxD = max(Dst(:));
    Cost = (Dst./maxD).^-4; Cost(Dst==0) = Inf;
    %temp = ~isinf(Cost) & SEED1 & SEED2; %check for connectivity
    clear Dst
    % Do fast marching using the maximum distance value in the image
    % and the points describing all found branches are sourcepoints.
    fprintf('Fast marching & skeletonization...\n');
    tic
    GD1 = graydist(Cost,SEED1,'quasi-euclidean'); toc

    GD2 = graydist(Cost,SEED2d,'quasi-euclidean'); toc

    
    GDm = GD1+GD2;
    S = GDm<(min(GDm(:))+0.1);
    tic; S2 = bwskel(S); toc
    %if the points are in the corners, we may end up with artifacts going along
    %the edges. Remove those by clearing the borders:
    S2(1,:,:) = false; S2(end,:,:) = false; S2(:,1,:) = false; S2(:,end,:) = false;
               
    clear S

    fprintf('Saving skeletonization variables...\n');
    save([fileLoc '\' IM_FILE(1:end-4) '_skmSKEL.mat'],'S2','GD1','GD2','GDm','Cost','-v7.3');
end

clear GD1 GD2 GDm Cost

%ok now we need to find the direction of the muscle along the skeleton
%brute force is to visit each point and calculate local direction.
fprintf('Calculate local direction...\n');
tic
EP = bwmorph3(S2,'endpoints'); %find endpoints
ind = find(EP,1,'first'); %we only need one endpoint
EP = false(size(EP)); EP(ind) = true;
Dleng = bwdistgeodesic(S2,EP,'quasi-euclidean'); toc

pts = find(S2);
nhood = ones(51,51,51);
MuscDirAll = zeros(size(pts,1),3);
tic
for ip = 1:length(pts)
    val = Dleng(pts(ip));
    TEMP = Dleng>(val-25.5) & Dleng<(val+25.5);
    if ~any(TEMP(:)) 
        TEMP(pts(ip)) = true;
        TEMP = imdilate(TEMP,nhood) & S2;
    end
    s = regionprops3(TEMP,'EigenVectors');
    MuscDirAll(ip,1:3) = s.EigenVectors{1}(:,1)';
end
toc

%add in the parallel/anti-parallel fix
%find the angle between the local vectors and the muscle vector
mlth = zeros(1,size(MuscDirAll,1));
for i = 1:size(MuscDirAll,1)
    C = dot(MuscDirVolume,MuscDirAll(i,:)); %dot product
    am = norm(MuscDirVolume); % should be 1
    bm = norm(MuscDirAll(i,:)); % should be 1
    
    mlth(i) = acosd(C/(am*bm));

end

%find the angle between the local anti-vectors and the muscle vector
mlnth = zeros(1,size(MuscDirAll,1));
for i = 1:size(MuscDirAll,1)
    C = dot(MuscDirVolume,-MuscDirAll(i,:)); %dot product
    am = norm(MuscDirVolume); % should be 1
    bm = norm(MuscDirAll(i,:)); % should be 1
    mlnth(i) = acosd(C/(am*bm));
end
%we don't actually care about parallel or antiparallel vectors, so use the
%smallest angle
swaplist = mlnth<mlth;
MuscDirAllNew = MuscDirAll;
MuscDirAllNew(swaplist,:) = MuscDirAllNew(swaplist,:)*-1;
MuscDirAll = MuscDirAllNew;

%ok, now we need to find the closest pt to each nuclei to find local muscle
%direction. Since the number of skeleton pts is relatively small, can brute
%force the distance measurement
MuscDirLocal = zeros(NumNuc,3);
[rp cp sp] = ind2sub(size(S2),pts);
for in = 1:NumNuc
    AllD = sqrt( (cx(in)-cp).^2 + (cy(in)-rp).^2 +  (cz(in)-sp).^2);
    Minpts = find(AllD==min(AllD));
    %demo closest point finding, easiest to do if you step faster to NumNuc
    %eg use 1:10:NumNuc for example
    %    figure; imagesc(max(S2,[],3)); hold on; plot(cy(in),cx(in),'go'); plot(cp(Minpts),rp(Minpts),'gx'); axis equal; pause(2); close(gcf)
    %average multiple points if needed
    MuscDirTemp = MuscDirAll(Minpts,1:3);
    MuscDirLocal(in,1:3) = mean(MuscDirTemp,1);
end


% figure(1)
% quiver3(X,X,X,V,U,W); hold on %swap U and V
% quiver3(0,0,0,MuscDirVolume(2),MuscDirVolume(1),MuscDirVolume(3),5)

%find the angle between the nuclei vectors and the muscle vector
mnth = zeros(1,size(NucDir,1));
for i = 1:size(NucDir,1)
    MuscDir = MuscDirLocal(i,1:3);
    C = dot(MuscDir,NucDir(i,:)); %dot product
    am = norm(MuscDir); % should be 1
    bm = norm(NucDir(i,:)); % should be 1
    mnth(i) = acosd(C);

    
end
%find the angle between the nuclei anti-vectors and the muscle vector
mnnth = zeros(1,size(NucDir,1));
for i = 1:size(NucDir,1)
    MuscDir = MuscDirLocal(i,1:3);
    C = dot(MuscDir,-NucDir(i,:)); %dot product
    am = norm(MuscDir); % should be 1
    bm = norm(NucDir(i,:)); % should be 1
    mnnth(i) = acosd(C);
 end

%testing you can pick a nuc vector and see if the angle looks right
% figure(2)
% subplot(2,1,1)
% numt = 500;
% mnth(numt)
% mnnth(numt)
% quiver3(0,0,0,MuscDirVolume(2),MuscDirVolume(1),MuscDirVolume(3)); hold on
% quiver3([0],[0],[0],NucStats(numt).EigenVectors(2,1),NucStats(numt).EigenVectors(1,1),NucStats(numt).EigenVectors(3,1)); axis equal
% quiver3([0],[0],[0],-NucStats(numt).EigenVectors(2,1),-NucStats(numt).EigenVectors(1,1),-NucStats(numt).EigenVectors(3,1));
% subplot(2,1,2)
% numt = 501;
% mnth(numt)
% mnnth(numt)
% quiver3(0,0,0,MuscDirVolume(2),MuscDirVolume(1),MuscDirVolume(3)); hold on
% quiver3([0],[0],[0],NucStats(numt).EigenVectors(2,1),NucStats(numt).EigenVectors(1,1),NucStats(numt).EigenVectors(3,1)); axis equal
% quiver3([0],[0],[0],-NucStats(numt).EigenVectors(2,1),-NucStats(numt).EigenVectors(1,1),-NucStats(numt).EigenVectors(3,1));

%we don't actually care about parallel or antiparallel vectors, so use the
%smallest angle
swaplist = mnnth<mnth;
NucDirNew = NucDir;
NucDirNew(swaplist,:) = NucDirNew(swaplist,:)*-1;

%we don't need to rederive the angles, we can just use the min of mnth and
%mnnth, but easy enough:
NDth = zeros(1,size(NucDirNew,1));
for i = 1:size(NucDirNew,1)
    MuscDir = MuscDirLocal(i,1:3);
    C = dot(MuscDir,NucDirNew(i,:)); %dot product
    am = norm(MuscDir); % should be 1
    bm = norm(NucDirNew(i,:)); % should be 1
    NDth(i) = acosd(C);
       tf = isreal(mnth(i));
   
end

% U = NucDirNew(:,1);
% V = NucDirNew(:,2);
% W = NucDirNew(:,3);
% 
% figure(3)
% quiver3(X,X,X,V,U,W); hold on %swap U and V
% quiver3(0,0,0,MuscDirVolume(2),MuscDirVolume(1),MuscDirVolume(3),5)


%testing for things pointing the right direction
%create labeled subset of nuclei
% NumNuc = size(NucDirNew,1);
% R = randperm(NumNuc); R = R(1:100); %randomly select x nuclei...you can select them all it's just hard to see
% Nucsub = ismember(L, R);
% tic
% figure(4)
% [faces,verts] = isosurface(Nucsub,0.5);
% colors = zeros(length(verts),3); colors(:,1) = 1;
% patch('Vertices',verts,'Faces',faces,'FaceVertexCData',colors,...
%     'FaceColor','interp','EdgeColor','interp')
% view(3) ; hold on;
% quiver3(cx(R),cy(R),cz(R),V(R),U(R),W(R),0.5); hold on %swap U and V
% quiver3(1000,1000,60,MuscDir(2),MuscDir(1),MuscDir(3),5)
% toc
% 
% figure(5)
% Uml = MuscDirAll(:,1);
% Vml = MuscDirAll(:,2);
% Wml = MuscDirAll(:,3);
% quiver3(cp(1:20:end),rp(1:20:end),sp(1:20:end),Vml(1:20:end),Uml(1:20:end),Wml(1:20:end),0.5); axis equal
% 
% 
% figure(6)
% U = MuscDirAll(:,1);
% V = MuscDirAll(:,2);
% W = MuscDirAll(:,3);
% 
% quiver3(zeros(size(U)),zeros(size(U)),zeros(size(U)),V,U,W); hold on %swap U and V
% quiver3(0,0,0,MuscDirVolume(2),MuscDirVolume(1),MuscDirVolume(3),5)

fprintf('Saving data variables...\n');
save([fileLoc '\' IM_FILE(1:end-4) '_DATA2.mat'],'s','NucStats','NDth','MuscDirAll','-v7.3');


% %testing for things pointing the right direction
% %create labeled subset of nuclei
% NumNuc = size(NucDirNew,1);
% R = randperm(NumNuc); R = R(1:100); %randomly select x nuclei...you can select them all it's just hard to see
% Nucsub = ismember(L, R);
% tic
% figure(4)
% [faces,verts] = isosurface(Nucsub,0.5);
% colors = zeros(length(verts),3); colors(:,1) = 1;
% patch('Vertices',verts,'Faces',faces,'FaceVertexCData',colors,...
%     'FaceColor','interp','EdgeColor','interp')
% view(3) ; hold on;
% quiver3(cx(R),cy(R),cz(R),V(R),U(R),W(R),0.5); hold on %swap U and V
% quiver3(1000,1000,60,MuscDir(2),MuscDir(1),MuscDir(3),5)
% toc
```
