## Supplementary Material; VBAM Nuclear Alignment and Aspect Ratio Code for "*In vitro* formation and extended culture of highly metabolically active and contractile tissues": VBAM_NucAlignmentQuant_shell_EXAMPLE_v2023july5.m

```
%%% use with quant script (VBAM_NucAlignmentQuant_EXAMPLE_v2023july5)
%%
%Authors: Isabella A. Bagdasarian, Joshua T. Morgan
%Lab: TIME lab, PI:Dr. Joshua Morgan, Bioengineering Department
%Institution: Univerisity of California, Riverside
%last edited 07/05/2023 IAB

% Description: Prepares output directories and organizes .lif file
% directory for downstream processing with
% VBAM_NucAlignmentQuant_EXAMPLE_v2023july5. Allows for user defined flags
% to redo chunks of analysis as needed. Allows for users to specify data
% input directory (dataLoc) and output directory (fileLoc).

clearvars
close all
clc

%% FLAGS & USER INPUTS

REDO_stitch = 0; %set to 1 if stitching needs to be redone
REDO_nuc = 0; %set to 1 if nuclei seg needs to be redone
REDO_WS = 0; %set to 1 if nuclei watershedding needs to be redone
REDO_skm = 0; %set to 1 if nuclei watershedding needs to be redone

% specify data location and save location
dataLoc = 'H:\DenseCultures_VBAMCellularity_lifs\';

% offset storage location for stitching
offsetLoc = 'D:\Bella\Muscle\offsets\';

%image output
outdir1 = 'H:\Bella\Muscle\analysis\';
namey = 'NucAlignment Quant';
outdir2 = ['output' '_' namey '_' '31-March-2023' '\'];%define saving folder


fileLoc = [outdir1 outdir2]; %define full saving location
if ~isdir(sprintf('%s',fileLoc))%make a folder if it doesn't exist
    mkdir(sprintf('%s',fileLoc))
end

%% make file list
FileList1 = dir([dataLoc, '*.lif']);
lg = (length(FileList1));

%% 
for file = [1:lg]
    Stic = tic;
    IM_FILE = FileList1(file).name;
    
    BaseFileName = sprintf('%s', IM_FILE(1:end-4));
    
    fprintf(sprintf('Begin analysis on %s... \n',BaseFileName))
    
    VBAM_NucAlignmentQuant_EXAMPLE_v2023july5
    
    Stoc = toc(Stic);   fprintf(1,'Analysis took %f seconds.\n',Stoc)

    clearvars -except lg FileList1 fileLoc outdir1 outdir2 ...
        offsetLoc dataLoc REDO_stitch REDO_skm REDO_nuc REDO_WS ...
    
 end
```
