## Supplementary Material; VBAM Nuclear Alignment and Aspect Ratio Code for "*In vitro* formation and extended culture of highly metabolically active and contractile tissues": VBAM_NucAspectRatio_dataViz_EXAMPLE_v2023july5.m

```
clearvars
close all
clc

%load output NucStats data from VBAM_NucAlignmentQuant
dataVars = dir(['D:\Bella\Muscle\analysis\output_NucAlignment Quant_20-February-2023\data outputs\', '*.mat']);

% FIND NUCLEI ASPECT RATIO
for j = 1:length(dataVars)
    
    clear NucStats;
    BaseFileName = dataVars(j).name;
    BaseFileName = BaseFileName(1:end-4);
    load(sprintf('%s\\data outputs\\%s','D:\Bella\Muscle\analysis\output_NucAlignment Quant_20-February-2023',dataVars(j).name),'NucStats');
    
    axisL = vertcat(NucStats.PrincipalAxisLength); %extarct principal axis length
    
    aspR = axisL(:,1)./axisL(:,2); %divide major axis by minor axis
    
    allAspR{j} = aspR'; %collect nuclei aspect ratio data into single variable
end
```
