## Supplementary material for "*In vitro* formation and extended culture of highly metabolically active and contractile tissues": Table 2

| **Primary Antibody** | **Source** | **Concentration** | **Use** |
|---|---|---|---|
| *DyLight 554 Phalloidin* | 13054; Cell Signaling Technologies, Danvers, MA | [1:200] | F-actin marker |
| *DRAQ7* | 7406; Cell Signaling Technologies | [1:100] | Nuclear marker |
| *Myosin (Fast) mouse monoclonal* | M1570; Sigma-Aldrich | [1:250] | Muscle differentiation marker[1,2] |
| *Titin mouse monoclonal* | 9 D10; Developmental Studies Hybridoma Bank, Iowa City, IA | [1:100] | Muscle differentiation marker[3,4] |
| *mCherry rabbit polyclonal* | PA5-34974; Invitrogen, Rockford, IL | [1:250] | Label mCherry tagged HMEC1 cells |
| **Secondary Antibody** | **Source** | **Concentration** | **Use** |
| *Goat Anti-Rabbit IgG DyLight™ 550 Conjugated* | 84541; Invitrogen | [1:500] | mCherry secondary |
| *Goat Anti-Mouse IgG, DyLight™ 488 Conjugated* | 35502; Invitrogen | [1:500] | Myosin or Titin secondary |

| **BLOCKING BUFFER** (500 mL) | | |
|---|---|---|
| ***Reagent*** | ***Source*** | ***Amount*** |
| $ddH_2O$ | - | 450 mL |
| 10 x PBS | Apex Bioresearch Products, Houston, TX #20-134 | 50 mL |
| Bovine Serum Albumin (BSA) | Genesee Scientific 25-529 | 5 g |
| Tween 20 | Fisher BioReagents BP337 | 0.5 mL |
| Cold water Fish Gelatin | Sigma-Aldrich G7041 | 1 g |
| Sodium Azide (10% Sodium Azide in $diH_2O$) | Fisher Chemical S227I | 5 mL (0.1% final concentration) |
