## Supplementary Figures for "*In vitro* formation and extended culture of highly metabolically active and contractile tissues"

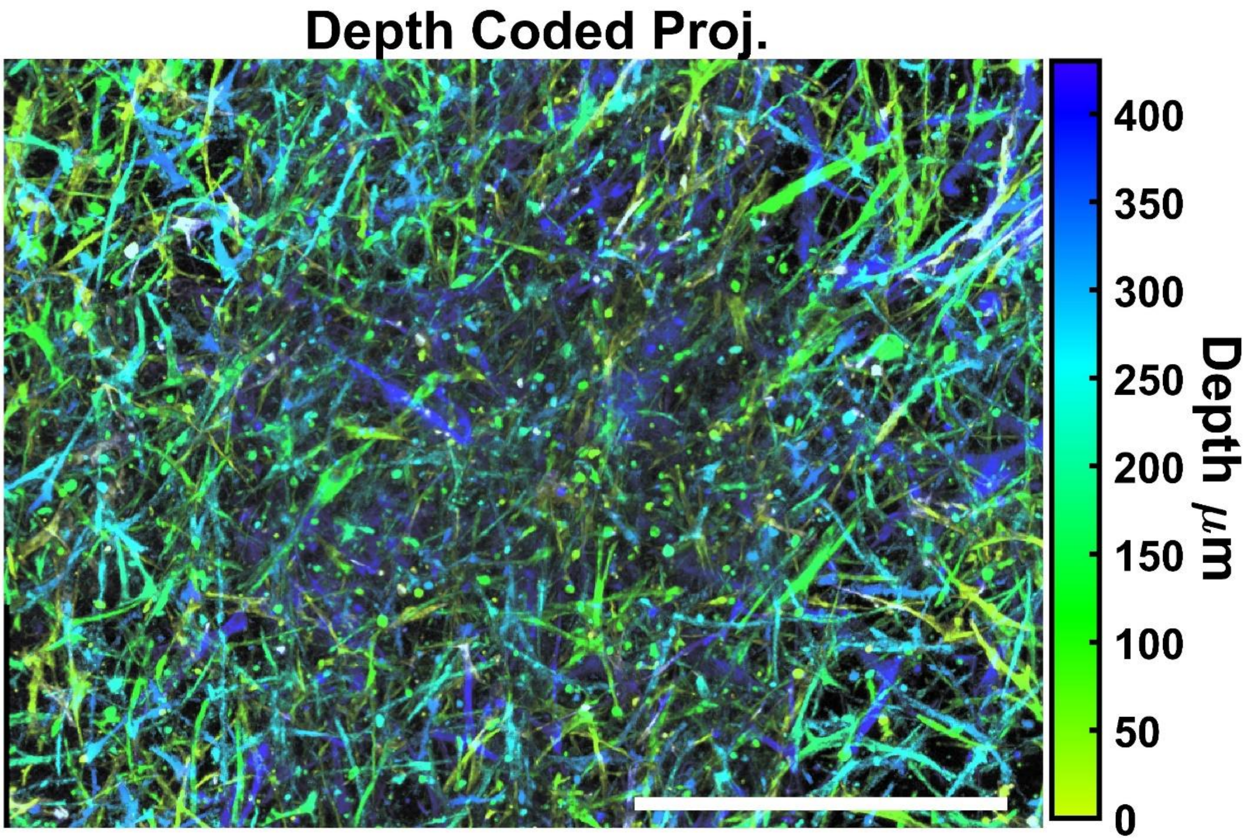


**Figure S1**. VBAM differentiation media supports vascular network formation in EC/ASC co-culture. ECs and ASCs were cultured in VBAM differentiation media with VEGF (weeks 0-2) or PDGF-BB (weeks 2-4) before being fixed and stained against collagen IV (EC basement membrane marker). Shown is a depth-coded projection of a stitched tilescan demonstrating vascular networks have assembled and are present through the culture bulk (scalebar = 250 μm).

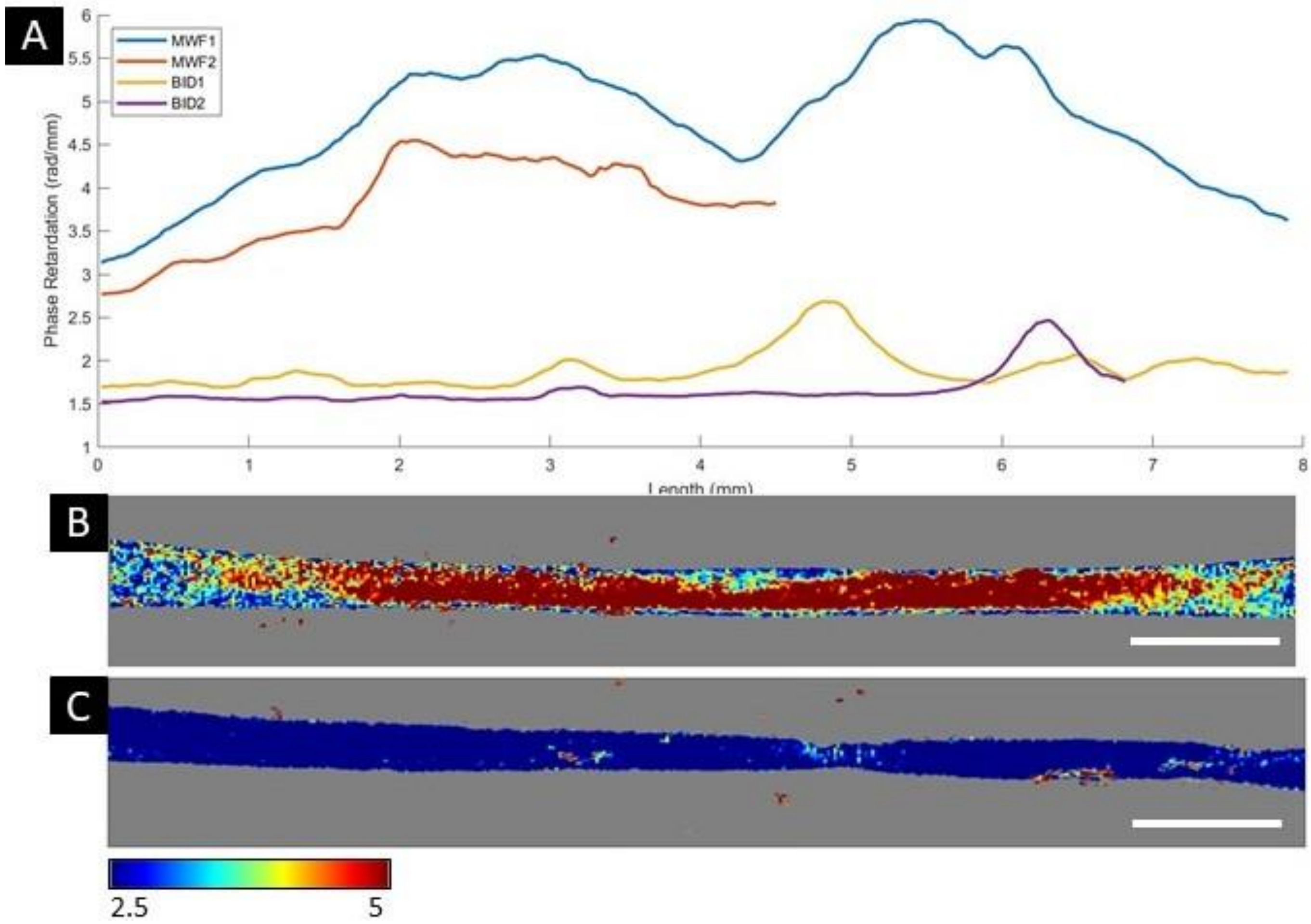


**Figure S2.** (A) The average phase retardation as a function of sample length for the MWF and B.I.D samples. The whole length of the B.I.D2 sample could not be obtained because of the pins used to fix the samples during imaging. (B-C) Representative en face phase retardation images (in rad/mm) taken from the middle of the MWF1 and BID1, respectively (scalebar = 1 mm). The MWF samples have higher phase retardation throughout the length of the samples compared to the B.I.D samples.

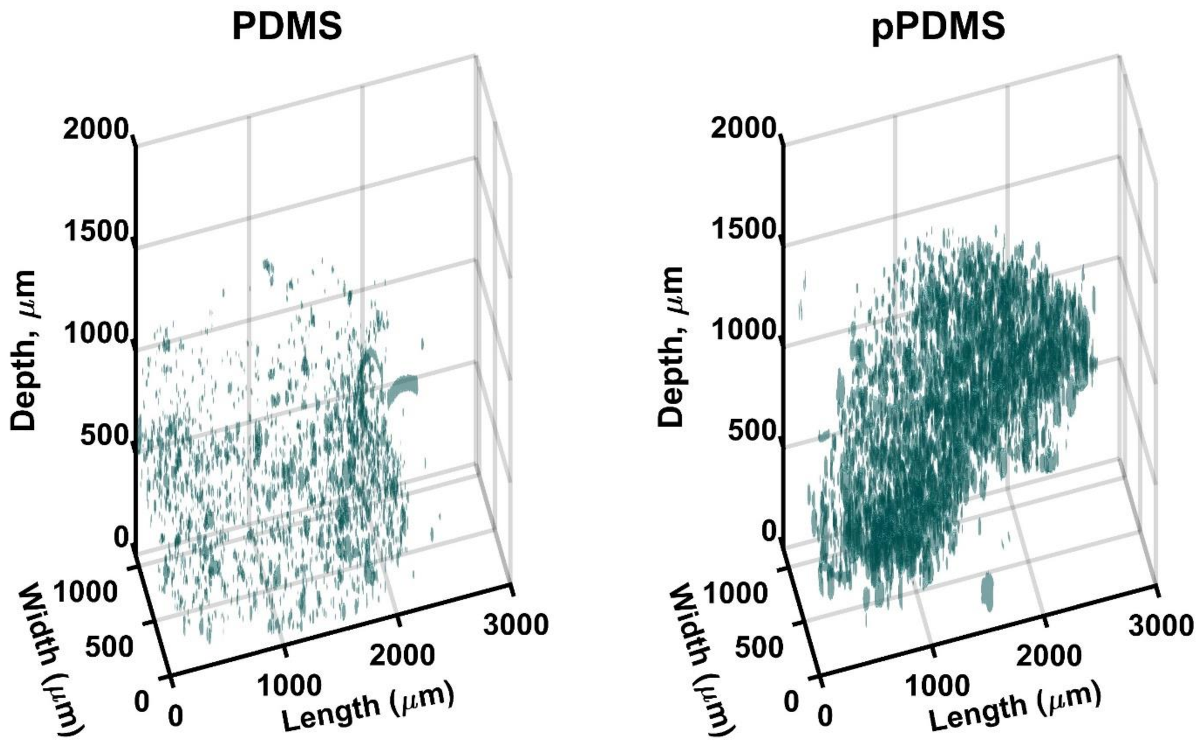


**Figure S3.** 3D renderings of segmented nuclei of S-PDMS and P-PDMS cultures maintained via side wall diffusion. Stitched tilescans were rendered in 3D to visualize spatial distribution of nuclei. Density of P-PDMS nuclei appears increased with evidence of dense aggregate formation relative to S-PDMS.

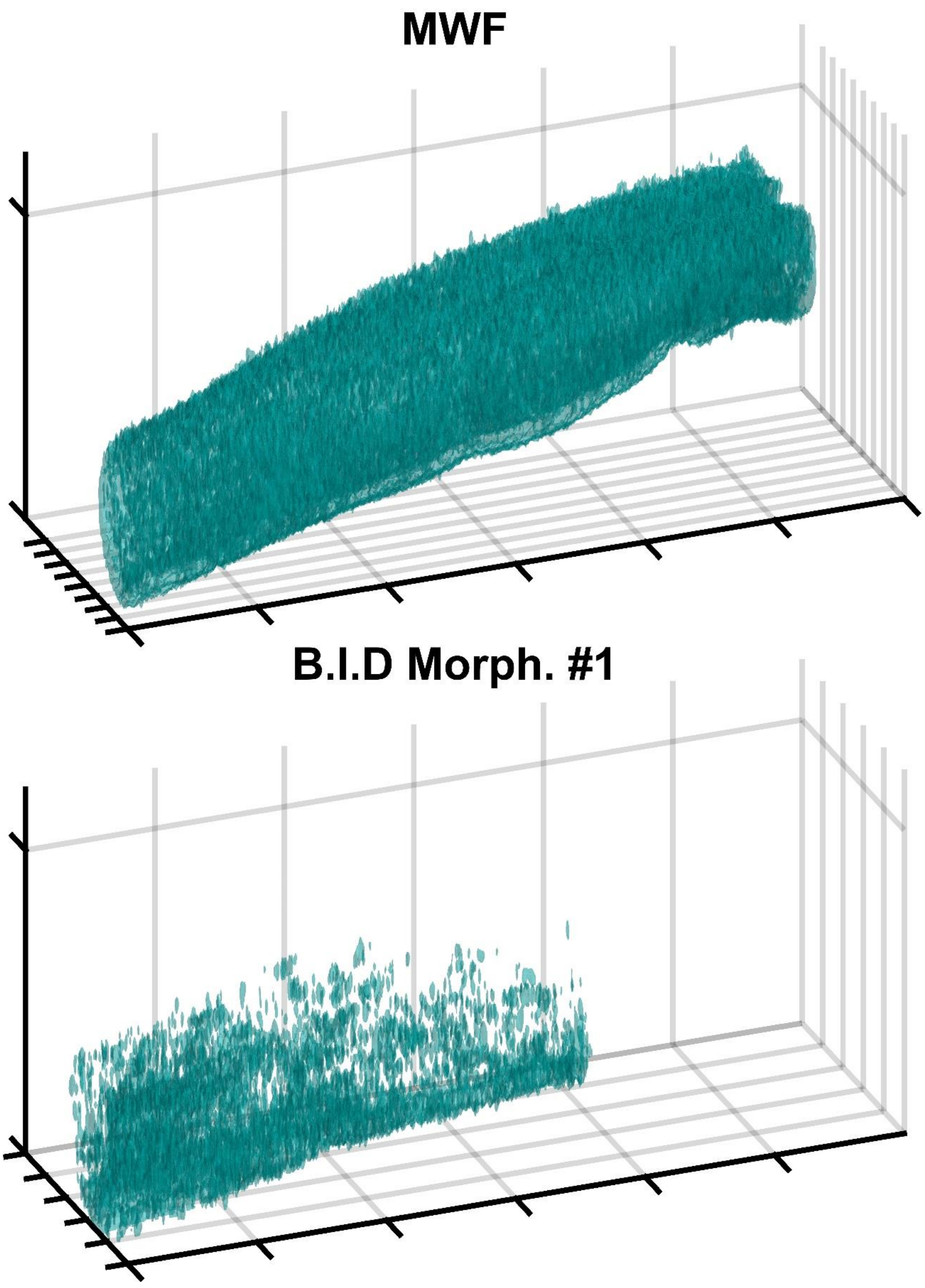


**Figure S4.** 3D renderings of segmented nuclei in stitched tilescans of MWF and B.I.D Morph #1 (3 mm x 1 mm x 0.6 mm). Nuclei density is increased with consistent organization globally in MWF VBAMs.

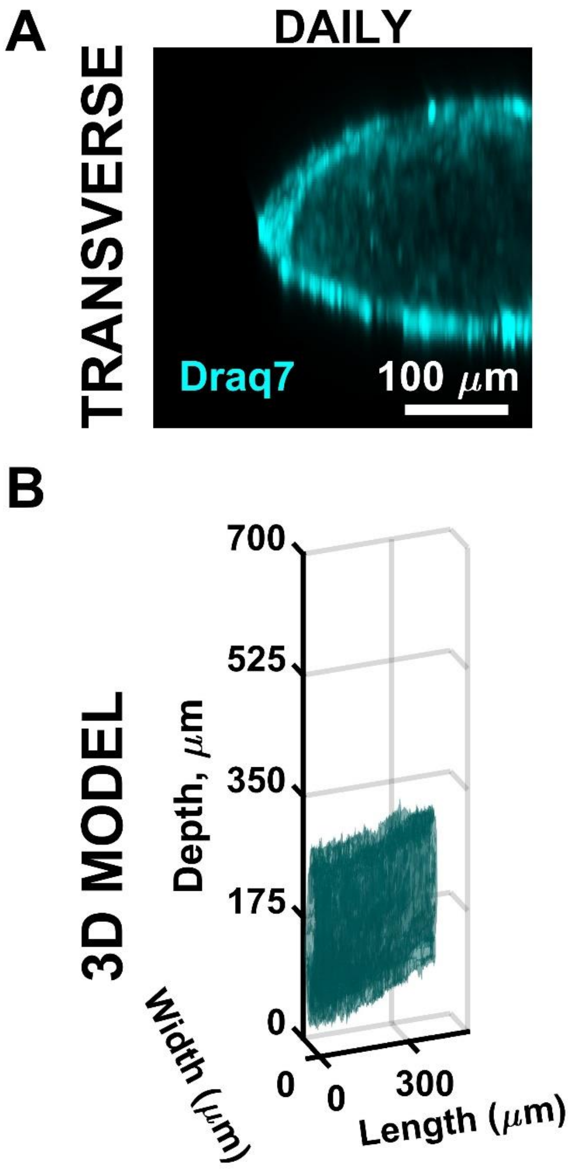


**Figure S5.** Effects of daily feeding on VBAM cellularity. (A) Representative transverse projection of VBAM nuclei after 5 weeks of daily media changes show similar nuclei density to MWF samples. (B) 3D rendering of segmented nuclei of daily fed VBAM indicate similar spatial organization of the nuclei to MWF samples.

**Figure Legends**

**Figure S1**: VBAM differentiation media supports vascular network formation in EC/ASC co-culture. ECs and ASCs were cultured in VBAM differentiation media with VEGF (weeks 0-2) or PDGF-BB (weeks 2-4) before being fixed and stained against collagen IV (EC basement membrane marker). Shown is a depth-coded projection of a stitched tilescan demonstrating vascular networks have assembled and are present through the culture bulk (scalebar = 250 µm).

**Figure S2**: (A) The average phase retardation as a function of sample length for the MWF and B.I.D samples. The whole length of the B.I.D2 sample could not be obtained because of the pins used to fix the samples during imaging. (B-C) Representative en face phase retardation images (in rad/mm) taken from the middle of the MWF1 and BID1, respectively (scalebar = 1 mm). The MWF samples have higher phase retardation throughout the length of the samples compared to the B.I.D samples.

**Figure S3:** 3D renderings of segmented nuclei of S-PDMS and P-PDMS cultures maintained via side wall diffusion. Stitched tilescans were rendered in 3D to visualize spatial distribution of nuclei. Density of P-PDMS nuclei appears increased with evidence of dense aggregate formation relative to S-PDMS.

**Figure S4**: 3D renderings of segmented nuclei in stitched tilescans of MWF and B.I.D Morph #1 (3 mm x 1 mm x 0.6 mm). Nuclei density is increased with consistent organization globally in MWF VBAMs.

**Figure S5:** Effects of daily feeding on VBAM cellularity. (A) Representative transverse projection of VBAM nuclei after 5 weeks of daily media changes show similar nuclei density to MWF samples. (B) 3D rendering of segmented nuclei of daily fed VBAM indicate similar spatial organization of the nuclei to MWF samples.
