## Supplementary material for "*In vitro* formation and extended culture of highly metabolically active and contractile tissues": Table 1

| Media | Source | Components |
|---|---|---|
| IMR90 Fibroblast cell line | Corning, Manassas VA #10-090-CV | DMEM/F12 50:50 |
| | Corning #35-010-CV | 10% Fetal Bovine Serum (FBS) |
| | Genesee Scientific, El Cajon, CA #25-512 | 1% Penicillin/Streptomycin (P/S) |
| MDCK Epithelial cell line | Corning #10-013-CM | DMEM HG |
| | * | 3% FBS |
| | * | 1% P/S |
| | VWR Life Sciences, Radnor, PA #IC0219504391 | Amphotericin B [250 µg/mL] |
| HMEC-1 Endothelial cell line | Gen Depot, Baker, TX #CM034-050 | MCDB131 Base medium |
| | * | 10% FBS |
| | * | 1% P/S |
| | Fisher Bioreagents, Waltham, MA #BP379-100 | L-Glutamine [10 mM] |
| | Peprotech, Cranbury, NJ #AF-100-15-1MG | Epidermal Growth Factor (EGF) [10 ng/mL] |
| | Alfa Aesar, Tewksbury, MA A1629203 | Hydrocortisone [10 µg/mL] |
| | * | Amphotericin B [250 µg/mL] |
| C2C12 Myoblast cell line | Corning #10017CM | DMEM HG without Sodium Pyruvate |
| | * | 10% FBS |
| | * | 1% P/S |
| | * | Amphotericin B [250 µg/mL] |
| ASC52telo adipose derived stem cell line | ATCC, Manassas VA #PCS-500-030 | Mesenchymal Stem Cell (MSC) Basal Medium |
| | ATCC #PCS-500-030 | 2% FBS |
| | ATCC #PCS-500-030 | FGF basic [5 ng/mL] |
| | ATCC #PCS-500-030 | FGF acidic [5ng/mL] |
| | ATCC #PCS-500-030 | L-Alanyl-L-Glutamine [2.4 mM] |
| | ATCC #PCS-500-030 | G418 [0.2 mg/mL] |
| VBAM growth media (day 0-4 of culture) | * | MCDB131 Base medium |
| | * | 10% FBS |
| | * | 1% P/S |
| | * | Amphotericin B [250 µg/mL] |
| | * | L-Glutamine [10 mM] |
| | * | Epidermal Growth Factor (EGF) [10 ng/mL] |
| | * | Hydrocortisone [10 µg/mL] |

| | | |
|---|---|---|
| | Fisher Chemical, Fair Lawn, NJ #A61-100 | L-Ascorbic Acid [50 μg/mL] |
| | TCI, Portland, OR #H0296 | Trans-4-Hydroxy-L-proline [10 mg/L] |
| | Fisher Bioreagents #BP392-100 | L-Proline [40 mg/L] |
| | Peprotech 100-20-1MG | VEGF [1 ng/mL] |
| VBAM differentiation media (day 5-40 of culture) | * | MCDB131 Base medium |
| | * | 2% FBS |
| | * | 1% P/S |
| | * | Amphotericin B [250 μg/mL] |
| | * | L-Glutamine [10 mM] |
| | * | Epidermal Growth Factor (EGF) [10 ng/mL] |
| | * | Hydrocortisone [10 μg/mL] |
| | Gibco, Grand Island, NY #41400045 | 1x Insulin-Transferrin-Selenium |
| | * | VEGF [1 ng/mL] (day 5-14) |
| | * | PDGF-BB [0.1 ng/mL] (day 14-40) |

*** Indicates source information was previously defined.**
